# Emergent Tissue Stability from Intercellular Bond Dynamics during Cyclic Mechanical Loading

**DOI:** 10.1101/2025.11.27.691035

**Authors:** Eleni Papafilippou, Alessandra Bonfanti, Guillaume Charras, Alexandre Kabla

## Abstract

Epithelial tissues are often exposed to cyclic deformations in their physiological environment. The mechanical integrity of epithelial tissues relies on intercellular adhesion proteins which link neighbouring cells and transmit forces across the cellular network. The stability of the intercellular adhesive bonds is critical. Under sustained stress, bond failure leads to cumulative damage that progressively weakens the material and ultimately causes failure at the tissue scale. Although the collective behaviour of adhesion bonds under static loading has been characterised to some extent, their response to dynamic loading remains largely unknown, despite its physiological importance. Here we combine quantitative experiments on MDCK monolayers with analytical and stochastic modeling of slip-bond intercellular linkers to investigate tissue resilience under cyclic loading. We find that cyclic loading significantly prolongs tissue lifetime and deformation before failure compared to constant tension, due to intervals of low stress facilitating bond repair. Our model identifies intrinsic rupture and repair timescales governing adhesion stability, revealing three regimes of tissue behavior: rupture, slow damage accumulation and stable equilibrium. Normalizing loading parameters by the intrinsic molecular timescales collapses experimental and simulated data into universal stability maps. These findings demonstrate that epithelial resilience emerges from stochastic adhesion bond dynamics, providing a predictive framework linking molecular adhesion turnover to macroscopic tissue mechanics under physiological cyclic forces.

**SIGNIFICANCE:** Epithelial tissues, the body’s primary mechanical barriers, experience continuous mechanical stress from physiological movements such as breathing or peristalsis. Through experiments and modeling, we show that a tissue’s ability to resist and repair damage under cyclic loading comes from the natural binding and unbinding of the molecules that link neighboring cells. We identify fundamental timescales governing rupture and recovery in cell-cell junctions, showing that tissue stability can be universally predicted from molecular adhesion kinetics. This work establishes a quantitative framework connecting microscopic adhesion kinetics to macroscopic tissue mechanics. It provides a tool for studying resilience in diverse biophysical soft matter systems and for understanding how living materials maintain integrity under fluctuating forces.

## INTRODUCTION

Tissue integrity and barrier strength are fundamental to organ health, yet epithelial tissues are continually challenged by a range of dynamic mechanical environments. Physiological processes such as breathing, peristalsis, and movement impose cyclic loads that vary over seconds, while growth, pregnancy, and tumor expansion generate slow, sustained loads lasting months or years (1–3). While moderate cyclic loading can enhance cytoskeletal organisation and intercellular adhesion (4), irregular or high-frequency oscillations can destabilise adhesion and compromise barrier function, contributing to diseases like irritable bowel syndrome (IBS) (5–8). Experiments and theory indicate that soft epithelial tissues often develop microlesions or cracks under constant loads (9). In some contexts, such defects are homeostatic, as seen in goblet-associated fractures (GAFs) of the gut epithelium, which represent regulated, localized remodeling (8). In other cases, microcracks can nucleate and propagate, leading to tissue-scale rupture (10). Together, these findings suggest that epithelial tissues exist in a delicate balance between load patterns and mechanical stability. This raises a central question: how do epithelial tissues maintain their integrity when exposed to periodic mechanical loads?

In epithelial tissues, cohesion is maintained by intercellular adhesion proteins that form dynamic links between cells (11). The binding and unbinding of these molecular linkers are governed by intrinsic timescales (5). A variety of cellular mechanisms, ranging from cytoskeletal remodelling to mechanotransductive and biochemical signalling, contribute to the regulation of intercellular junctions (12, 13). At the molecular scale, intercellular bonds can display a range of behaviours: slip bonds, which become weaker and more likely to break as force increases (14) and catch bonds, which are paradoxically stabilised by intermediate levels of force (15). Yet, it is challenging to disentangle whether epithelial resilience can be explained purely by the emergent kinetics of junctional bonds, without invoking active feedback and signaling pathways (16). To what extent, then, does the ability of a tissue to retain its integrity in the face of repeated cycles of loading depend on the emergent mechanisms of molecular turnover of intercellular proteins?

Our theoretical understanding of junctional mechanics largely stems from modelling efforts. Building on Bell’s seminal model of force-dependent bond dissociation (14), many studies have used the slip-bond framework to predict junctional integrity under constant or monotonically increasing loads. Multi-bond stochastic models have shown how parallel bonds rupture probabilistically, producing exponentially distributed lifetimes and collective failure events (17, 18). These models provide insight into how junction strength scales with bond number, how variability emerges from stochastic rupture events, and how catch-bond kinetics can help to stabilise intercellular junctions. However, most applications to date have either focused on zero-load equilibrium states, or on sustained loads and force ramps, where the probability of rebinding becomes negligible (10, 19, 20). In contrast, cyclic loading introduces alternating phases of high stress, during which bond rupture occurs, and phases of low stress, when rebinding is more likely to take place. Using cyclic loading, we can investigate whether periodic relief from loading enables recovery, thereby shaping epithelial resilience over longer timescales. More generally, how do molecular rupture and repair events determine junctional integrity, and what periodic loading characteristics define different regimes of epithelial stability?

Here, we use experiments and modelling to frame a novel approach to test tissue resilience and illustrate the subtle interactions between mechanical loading patterns and junction stability. We compare constant and cyclic stress experiments on Madin-Darby Canine Kidney (MDCK) epithelial monolayers and interpret the results with analytical and stochastic models to explore how intercellular junctions accumulate damage and repair over time. Under sustained high stress, we identify a characteristic timescale which sets the limit of the cell-cell junction’s lifetime. At lower stress levels, we uncover a distinct repair timescale, that captures how cell-cell bonds reform to restore junctional integrity. Together, these emergent timescales capture how tissues resist stress across a broad range of loading conditions. Building on this framework, we examine tissue behaviour under cyclic loading, where alternating phases of high and low tension establish a threshold between rupture and stable recovery. Our results reveal distinct regimes: rupture, slow damage accumulation, and full recovery, whose boundaries are determined not by absolute load amplitude, but by how the durations of force application compare to intrinsic rupture and repair timescales. This unifying framework demonstrates that epithelial resilience arises as an emergent property of adhesion bond kinetics, offering a tool for understanding how tissues withstand physiological forces over the lifetime of the organism.

## RESULTS

### Evidence for healing of intercellular junctions in epithelial monolayers under cyclic loading

As a baseline for understanding tissue failure under cyclic loading, we first consider the limit case of creep loading where the material is brought to and kep at a constant stress value. Previous studies have shown that epithelial sheets fail within a narrow range of tensions, corresponding to the mechanical limit of intercellular adhesion (10). We selected a load close to this range to challenge tissue integrity without inducing immediate catastrophic failure. We applied a step in stress implemented through PID-controlled feedback (Methods, 1.5) and quantified the load using a force transducer (Appendix 1). Using an *in vitro* model of MDCK monolayers (Methods, 1) suspended between the two rods of a custom-built stretching device (Figure 1a,b), previously described in (21). We applied constant tensions (Figure 1c) with a mean centered around 0.15 N, m^−1^ ± 0.05 N, m^−1^). The time to the first detectable cellular-scale defect within the tissue was defined as the rupture timescale t^∗^. For a given tension value, there is some stochastic variation in the rupture timescales across samples, but no evidence of systematic differences (see Appendix 2, Figure 8). Across a broad range of applied tensions (0.025 - 0.3 N, m^−1^), higher tension is associated with a faster characteristic rupture time, t^∗^ (Figure 1e). The range of rupture times under creep indicate that failure could occur stochastically even under identical sustained loads. The creep data therefore provides an average material timescale for a given tension value, beyond which epithelia cannot maintain cohesion. This observation raises a number of important questions: Is the molecular damage accumulated during constant loading reversible? Can rupture be delayed by intermittently relieving tension to allow molecular-scale repair?

**Figure 1:**
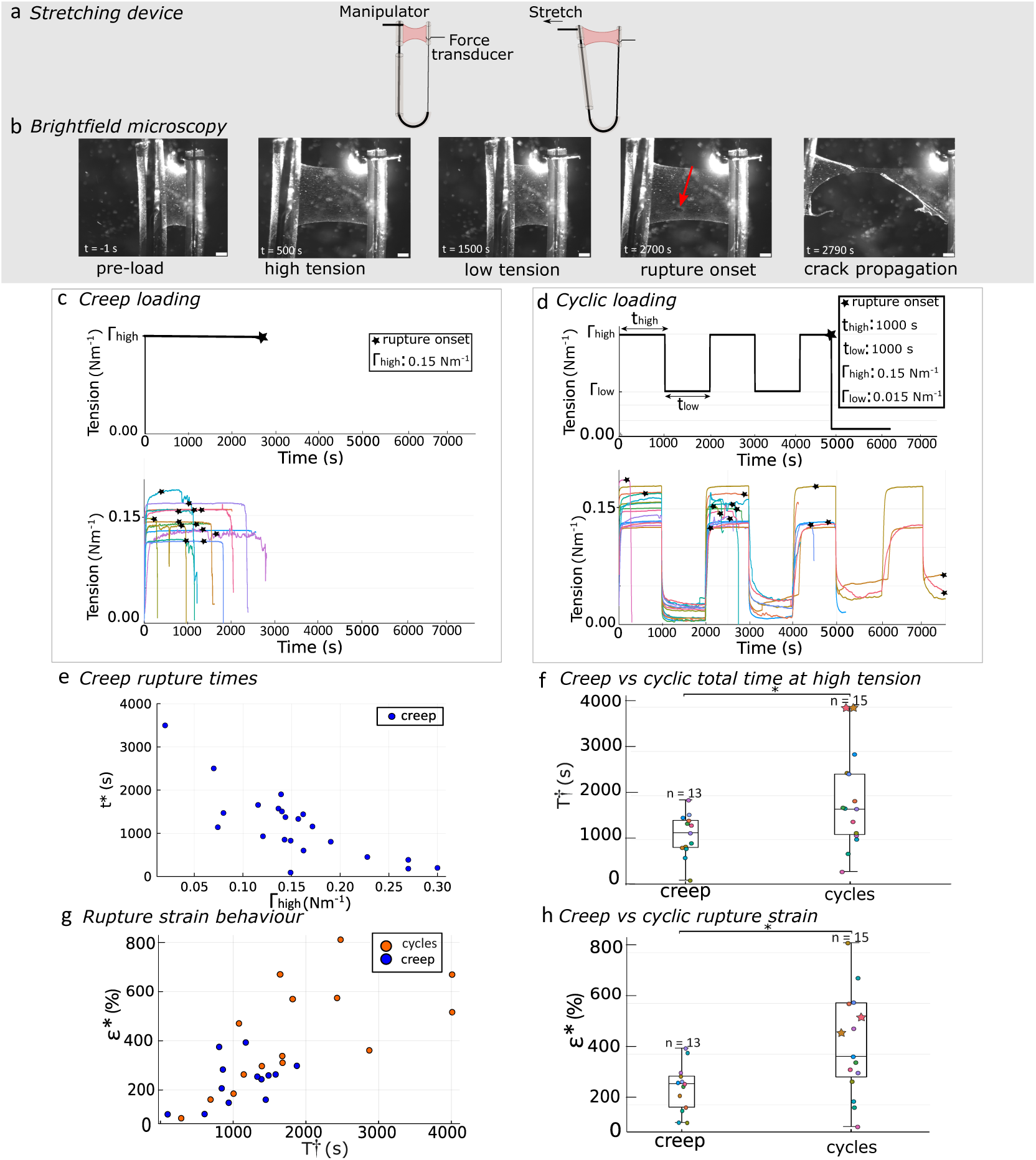
MDCK epithelial monolayers are more resilient to rupture when subjected to cyclic loading than under creep loading. a) Tissue stretching device consisting of a monolayer suspended between a thick wire attached to a motorised micromanipulator and a thin pliable wire hooked onto a force transducer. b) Bright-field microscopy time series of an MDCK monolayer during tension cycles. The rupture time, t^∗^, is defined as the time of the first defect and is indicated with a red arrow. Scale bars, 500*μ*m. c) Top: Idealised schematic of creep loading at tension Γ_high_. Bottom: Experimental creep loading data (moving average smoothing, window size 3.5s), curves of different colours represent different samples and the stars denotes rupture. d) Top: Idealised schematic of cyclic loading between high tension, Γ_high_ and low tension, Γ_low_. The durations of high and low tension are t_high_ and t_low_ respectively. Bottom: Experimental cyclic loading data (moving average smoothing, window size 3.5s) with Γ_high_ centred at 0.15Nm^−1^ and low tension, Γ_low_ centred at 0.015Nm^−1^. Both t_high_ and Γ_low_ durations are 1000s. Curves of different colours represent different samples and the stars denotes rupture. e) Creep rupture time t^∗^ for tension across two orders of magnitude. f) Box plots comparing the total time at Γ_high_, T^†^, under creep and cyclic loading. The central mark indicates the median, the top and bottom edges indicate the 25th and 75th percentiles respectively and the whiskers indicate the range excluding outliers. Stars indicate monolayers that did not break. *p=0.0382 for Wilcoxon rank sum test. g) Relationship between rupture strain, *ε*∗, and time at high tension, T^†^. h) Box plot comparing *ε*∗ for creep and cyclic loading, *p=0.0146 for Wilcoxon rank sum test.

To explore these questions, we subjected monolayers to cycles alternating periods of high and low tension. The high tension (Γ_high_) was applied for a duration t_high_ and the low tension (Γ_low_), was applied for a duration t_low_ (Figure 1d). The high tension target was 0.15 N.m^−1^ (± 0.05 N.m^−1^), matching the constant loading experiments to probe the onset of rupture. The target low tension was set to a small non-zero value, to ensure stable control by the PID feedback loop while maximizing the mechanical contrast between high and low tension phases. Experiments were limited to 8000s, beyond which cells progressively lose apico–basal polarity and monolayers start accumulating defects even without loading. Remarkably, MDCK epithelial monolayers showed, on average, enhanced mechanical resilience under cyclic loading compared to constant creep. When subjected to cyclic tension, monolayers maintained their mechanical integrity under high-tension states for longer (Figure 1f) relative to those subjected to constant creep. The total amount of time spent at high tension, T^†^, was compared across the two conditions and was found to be approximately 50% longer in the cyclic loading condition (creep mean lifetime: 1107s, cycles mean time spent at high tension: 1885s). A Wilcoxon rank sum test for independent samples, confirmed that this difference was statistically significant (*p=0.0382). Furthermore, five of the cyclic loading experiments, representing one third of all cyclic loading trials, exhibited markedly longer lifetimes when compared to the constant loading test data. Tukey’s HSD indicated these five observations as statistical outliers as they exceeded the 1.5 x IQR criterion for outlier detection. Notably, in two of these cases, the monolayer did not rupture within the experimental window (starred data points, Figure 1f), implying that their actual rupture times would extend even further. Thus, monolayers can endure high tension for substantially longer when subjected to cyclic loading compared to constant loading and the reported mean time spent at high tension for cyclic loading should be interpreted as a conservative lower bound.

In addition to rupture time, we compared the rupture strain under cyclic and monotonic loading to determine if rupture might be strain-controlled mechanism. Under constant stress, soft tissues typically display an initial regime of slow creep, followed by accelerated nonlinear deformation in which strain growth becomes increasingly rapid and ultimately diverges as the material approaches fracture (22). During cyclic loading, the global strain response reflects this progression, whereby after an initial slow creep regime, deformation increases nonlinearly and ultimately diverges as the monolayer approaches catastrophic failure (see Appendix 3, Figure 9). We quantified the strain in the monolayer and compared the strain at rupture, *ε*^∗^, across experimental conditions (Figure 1h). For both monotonic and cyclic loading, strain accumulated progressively during periods of applied tension, consistent with the viscoelastic nature of epithelial monolayers. Importantly, the rate of strain accumulation during high-tension phases was comparable across loading conditions (Figure 1g). As a result, cyclic loading, which on average maintained monolayers at high tension for longer durations, led to a greater cumulative deformation, *ε*∗, prior to rupture. So, even though the tissue experienced similar stress levels and stretched at the same rate, the monolayers that were stretched in cycles could deform more before rupture, indicating that failure is not determined solely by reaching a critical strain threshold. This observation argues against a strain-controlled failure mechanism. Interestingly, this also suggests that periodic unloading in the low-tension regime may allow the monolayer to reach greater cumulative deformation before failure. Introducing cycles appears to facilitate continued viscoelastic flow, highlighting that rupture occurs in a stress-limited regime. In such a regime, the sustained load is borne by cell-cell junctions, where stress concentration drives the gradual accumulation of damage (16, 23). To gain insight into the potential molecular origin of this phenomenon, we examine whether emergent junctional bond dynamics could underlie the delayed failure observed under cyclic loading conditions.

### Intrinsic timescales of molecular binding control the rupture and equilibrium dynamics of cell-cell junctions

A common approach to modelling intercellular protein dynamics is to use a slip bond, whose lifetime decreases exponentially with increasing force (Figure 2a) (14, 19, 24, 25). Bell’s classical model of receptor-ligand binding describes the force-lifetime relationship of a slip bond which can exist in one of two states: bound or unbound. The dynamics of a slip bond are governed by a force-dependent dissociation rate and a constant association rate (Equations 1a, 1b), where k_off,0_ is the dissociation rate constant in the absence of force, f is the externally applied force, and f_0_ is a characteristic force scale determined by the difference in free energy between the bound and unbound states (14). The application of tensile force lowers the energy barrier for bond rupture, thereby increasing the unbinding rate of individual slip bonds.

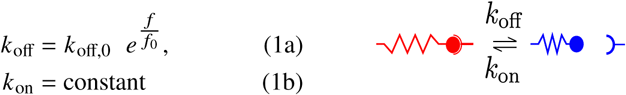

**Figure 2:**
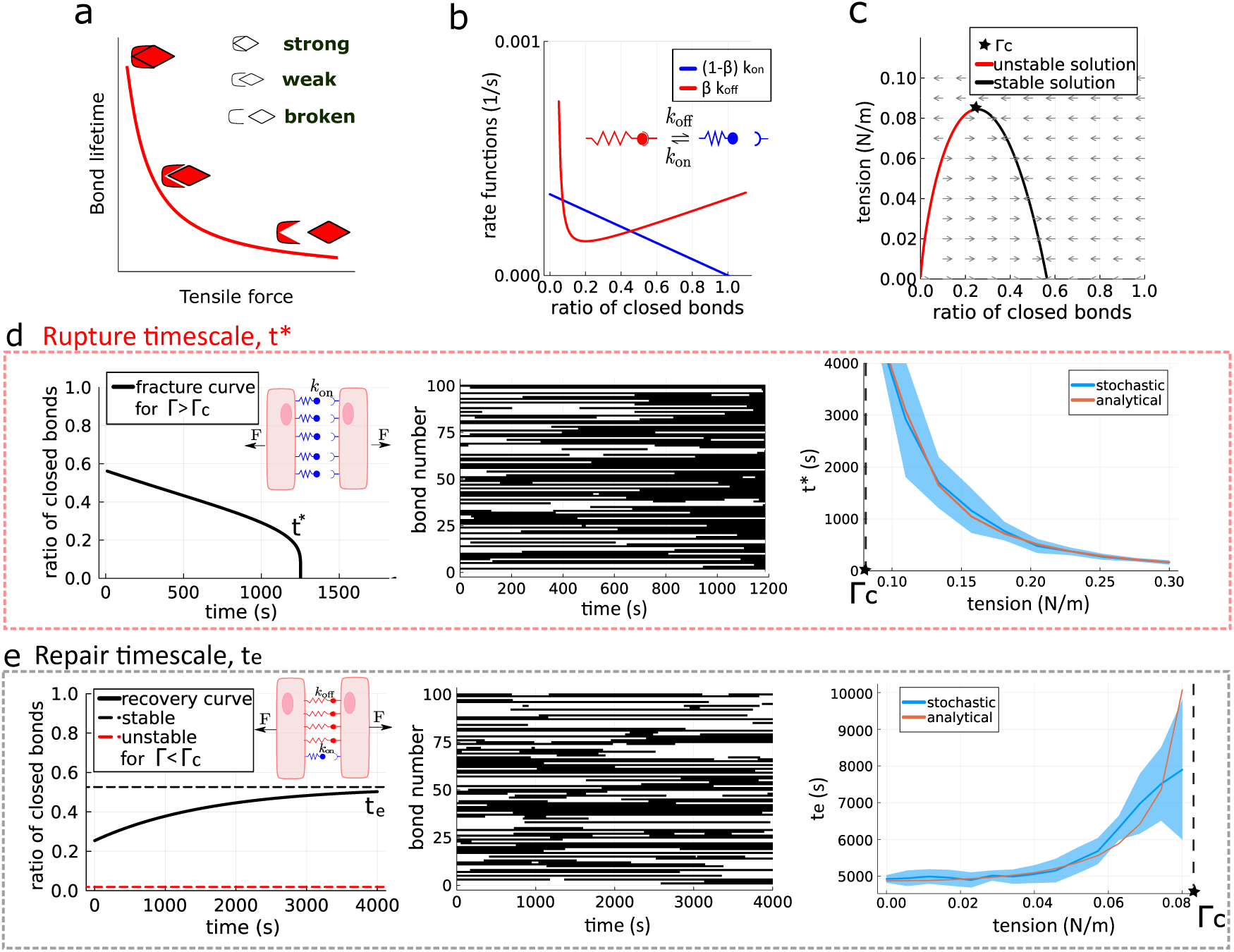
Theoretical framework for modelling an intercellular junction using a population of slip-bond linkers under force. a) A slip bond’s lifetime exponentially decays with increasing tensile force. b) Rate functions of link association (blue) and dissociation (red) as a function of the fraction of closed bonds, *β*, for a given tension (here, 0.04N/m). Points of equilibrium occur at the intersection between the two curves, where the rates of association and dissociation are equal. The stability of these points is determined by the local slopes of the two curves: a point is stable if small increases or decreases in the fraction of closed bonds produce net rates that drive the system back toward the intersection, and unstable if such deviations push it away. The values for the slip bond parameters, k_on_ and k_off,0_, are found in the Methods 2, Table 2. c) Quiver plot depicting stable (black) and unstable (red) analytical solutions for the ratios of closed bonds at different values of tension. The critical tension, Γ_c_, is the maximum tension value for which a solution exists. d) Determination of the rupture timescale *t*^∗^. Left: Rupture timescales, t^∗^, are defined for tension values larger than Γ_c_, initialised at the stable zero tension ratio of closed bonds. t^∗^ is calculated analytically by tracking the ratio of closed bonds over time. Middle: t^∗^ is calculated stochastically by tracking the states of 100 links over time (black is an open bond, white is a closed bond). Right: Analytical (orange) and stochastic (blue) rupture timescales as a function of applied tension decrease sharply with increasing tension and diverge near the critical tension Γ_c_. The blue shaded region represents the standard deviation of the stochastic model data. e) Determination of the repair timescale *t*_*e*_. Left: Repair timescales, t_e_, are defined for tension values smaller than Γ_c_, initialised at the stable ratio of closed bonds for a given tension value. t_e_ is calculated analytically by tracking the ratio of closed bonds over time. Middle: t_e_ is calculated stochastically by tracking the states of 100 links over time (black is an open bond, white is a closed bond). Right: Analytical (orange) and stochastic (blue) rupture timescales as a function of applied tension remain relatively constant at low tensions, but increase sharply as the applied tension approaches Γ_c_. The blue shaded region represents the standard deviation of the stochastic model data. Close to the transition point, there is an increase in standard deviation for the stochastic model’s t_e_.

Considering a population of N such linkers allows us to analytically describe how collective bond kinetics govern the stability of cell-cell junctions under mechanical stress. We define the fraction of closed linkers at any given time as *β*. The dynamics of *β* are governed by the balance between link association and dissociation (equation 2), where the first term represents the rate of link formation, which decreases linearly as *β* approaches unity, and the second term describes the rate of link dissociation, which increases exponentially with force. Γ denotes the applied tension (force per unit length) and 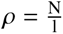 is the density of available linkers along a junction of length l. We assume tension is equally distributed across all closed links.

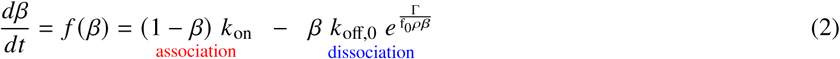

The rates of linker formation and linker loss can be written as functions of *β* (Figure 2b). At the intersection points of the two functions, the junction is in equilibrium, meaning that there is no net loss or gain of linkers over time, such that the rate of change of *β* is zero. The stability of each intersection point depends on the relative slopes of the two functions. For instance, if a small increase in *β* causes dissociation to dominate and a small decrease causes association to dominate, the system is driven back towards the intersection, making the equilibrium stable. In contrast, if deviations from the intersection are amplified, such that both an increase and decrease in *β* move the system away from the intersection point, then the fixed point is unstable. Increasing tension shifts the dissociation curve upward (in red, Fig 2b), which alters the position and eventually the number of intersection points. If there are no intersection points, there is no steady-state solution to equation 2 and therefore the cell-cell junction inevitably fails. Pairs of stable and unstable fixed points exist for tension values ranging from zero, up to a maximum tension, which we define as the critical tension value, Γ_c_ (Figure 2c). Conceptually, Γ_c_ represents the maximum mechanical load that an intercellular junction can bear. As such, Γ_c_ may also serve as a mechanical threshold for tissue integrity, beyond which stable cell adhesion cannot persist indefinitely.

The slip bond model parameters (see Methods, 2) were obtained by fitting the model’s fracture timescales to experimental fracture timescales using the experimental creep data. Further details on the fitting can be found Appendix 4, Figure 10). For convenience, we treat the product f_0_*ρ* as a single effective parameter, so that the influence of linker density is absorbed into an empirical fitting constant. The fitted parameters capture the experimental fracture dynamics well, with simulated fracture times closely matching the experimental measurements across the tested range (see Appendix 5, Figure 11). By varying model parameters such as the link kinetics or the applied tension, we can study how molecular changes affect the mechanical stability of intercellular junctions.

Our steady-state analysis identifies stable and unstable adhesion regimes determined by the applied tension. Having delineated these regimes, we then analyse how junction dynamics evolve towards them. We begin by considering the characteristic time to failure, which provides a direct comparison to the experimentally measured rupture time. For tension values larger than the critical tension Γ_c_, we define a rupture timescale, t^∗^, as the time required for all links to dissociate under constant tension, leading to junction failure (Figure 2d). For tension values below Γ_c_, we define a repair timescale, t_e_, as the time required for the junction to recover a stable equilibrium of bound linkers, under constant tension (Figure 2e), which can be calculated through the exponential half-life. Both timescales depend on the way the population is initialised; the rupture timescale was initialised at the zero-tension equilibrium fraction of closed bonds while the recovery timescale was initialised at the closed-bond fraction at Γ_c_. Further details can be found in Methods 2.

While the analytical model provides valuable insights into the average behaviour of adhesion at intercellular junctions, it inherently describes an idealised, deterministic system. In reality, adhesion proteins operate in a noisy environment with fluctuations that cannot be fully captured by a purely deterministic framework. Near the critical tension, stochastic behaviour can be the difference between a junction staying stable or failing catastrophically. To account for these effects, our analysis can be performed using a stochastic multi-bond framework governed by the same equations 1a and 1b. A cell-cell junction is defined as an interface with a number of links, N, which can be in one of two states: bound or unbound. The binding and unbinding of individual links with slip bond properties are treated as probabilistic events. The dynamics of a population of links can be simulated using a kinetic Monte Carlo scheme, initialised with a random distribution of states. We explored different values of N over three orders of magnitude (see Appendix 6, Figures 12,13) and chose N = 100, a value also used to capture MDCK monolayer dynamics and experimental noise levels in prior studies (10). For the stochastic model, t^∗^ was defined as the average time to failure across 100 creep simulations performed at high tension, each initialised at the zero-tension equilibrium fraction of closed bonds (see Methods 3). t_e_ was defined by tracking the states of 100 links over time as the average time to reach equilibrium under constant tension below Γ_c_, starting from the closed-bond fraction at Γ_c_. A further criterion was implemented, requiring that the starting point lie at least one standard deviation away from the stable fixed point, computed from the bond ratios across the 100 simulations at steady state. This criterion ensures that the repair timescale is clearly distinguished from stochastic fluctuations at steady state.

Rupture and repair timescales can be computed for a range of tensions. As tension increases from zero to Γ_c_, t_e_ increases exponentially, indicating slower recovery dynamics and increased variability near the transition point. In contrast, for tensions exceeding Γ_c_, t^∗^ decays exponentially, reflecting faster dissociation and mechanical failure under larger loads. When comparing the analytical and stochastic models, we find strong agreement across a wide tension range, reinforcing the idea that these transitions are governed by intrinsic timescales arising from bond-level kinetics. The main difference arises for the repair timescale t_e_ close to Γ_c_. For these tensions, a large number of stochastic trajectories fractured instead of repairing as reported in Appendix 7, Figure 15. Trajectories that ruptured were included only up to failure, to capture the initial recovery dynamics. This causes an underestimate of the analytical repair timescale close to the critical tension. An alternative definition of the stochastic repair timescales where trajectories that ruptured are assigned an infinite repair time is considered in Appendix 8. Therefore, the stochastic model of junctions predicts a gradual shift in the balance of rupture and repair probabilities, while the analytical model exhibits an absolute boundary between stable and unstable behaviour. In the following sections, we explore the interplay between the intrinsic timescales of bond dynamics and the extrinsic timescales of cyclic loads in tuning epithelial tissue behaviour.

### Intercellular junction stability under cyclic mechanical loads

We now investigate how cyclic loading affects the integrity of an intercellular junction. The applied load alternates between a high tension, Γ_high_, and a low tension, Γ_low_, sustained for durations of t_high_ and t_low_, respectively. For cyclic loading to meaningfully engage both the rupture and healing processes, Γ_high_ must exceed the critical tension Γ_c_, while Γ_low_ must remain below Γ_c_. To systematically explore the dynamic range of a junction’s response, we test a broad range of loading frequencies and duty ratios (the relative durations of t_high_ and t_high_ + t_low_) (Methods, 4). The results can be visualised through a map for the analytical (Figure 3a) and stochastic (Figure 3b) models. The colour bar quantifies the stability of the junction relative to the creep case, using the stability index, Θ, defined in equation 3. T^†^ is the total time spent at high tension until rupture (the same quantity previously used to analyse experimental data) and t^∗^ is the rupture timescale under constant high tension, as previously defined. For experimental data, we obtain Θ by averaging over independent repeats of an experimental condition, while for the stochastic model we average over 100 simulated trajectories. Average quantities are indicated by a bar in equation 3.

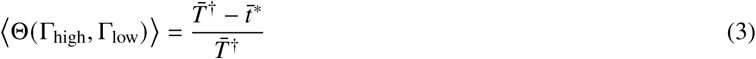

**Figure 3:**
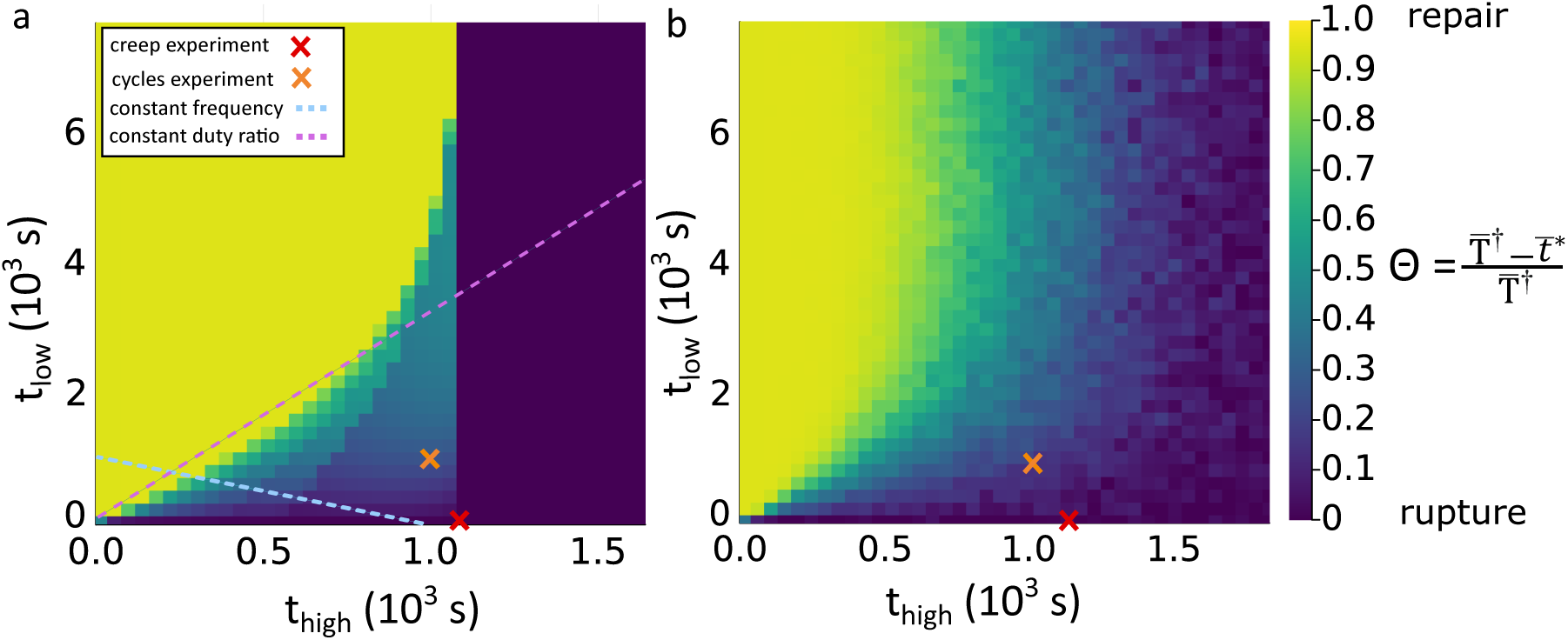
Models of intercellular linker dynamics reveal three distinct cell-cell junction behaviours in response to cyclic loading. a) Analytical heatmap illustrating junction behaviour under cyclic loading, defined between Γ_high_ = 0.15Nm^−1^ and Γ_low_ = 0.015Nm^−1^ (matching prior experimental conditions) for a broad range of combinations of t_high_ and t_low_. The colour scale encodes the normalised metric 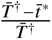 with values 1.0 (yellow) indicating full healing, 0 (purple) indicating immediate rupture and intermediate values representing a slow damage regime where linkers are progressively lost due to incomplete recovery during the low tension phase. The experimental data points can be overlaid on the map. The red cross indicates creep experiments leading to rupture at the critical time t^∗^ and the orange represents cyclic loading experiments (t_high_ = t_low_ = 1000s) that lie within the slow damage regime. The cyan dashed line denotes a constant frequency of 0.001Hz and the violet dashed line represents a constant duty ratio, 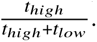 b) Stochastic heatmap defined with identical cyclic loading conditions.

When the time to rupture T^†^ is identical to the creep case (t_rupture_ = t^∗^), Θ = 0. When rupture doesn’t happen over the duration of the simulation, Θ = 1. Intermediate values (green-blue) are defined by the fractional delay in failure relative to t^∗^. Higher values indicate greater prolongation of junction lifetime compared to t^∗^, while lower values reflect only marginal extension. In this intermediate regime, junctional failure is not triggered by a single supercritical stress event, but by the progressive build-up of damage over many cycles. Each loading cycle gradually pushes the junction closer to rupture leading to a time to rupture T^†^ much longer than a single loading period. This regime reveals a fundamentally different pathway to failure that cannot be captured under monotonic or steady loading. Experimental observations from both creep and cyclic loading conditions can be mapped onto this phase diagram: creep-induced failure falls along the t_high_ axis and defines the t^∗^, Θ = 0, for a given tension (red cross Figure 3a), while cyclic loading resulted in delayed failure (orange cross, Figure 3a). Our analytical and stochastic models predicted values of Θ, close to one another and close to the experimentally observed value of Θ = 0.32 ± 0.27, an analytical Θ = 0.28 and a stochastic Θ = 0.23 ± 0.18.

Overall, our map reveals three distinct regimes of junction behaviour in response to cyclic loading: (i) stable repair, (ii) slow damage accumulation, and (iii) rupture. Moreover, the model demonstrates that transitions between regimes can be induced by tuning either the loading frequency or the duty ratio of t_high_ and t_low_ (Figure 3a). These observations raise several key questions: How can the emergent balance of linker loss and gain predict junctional stability? And how do changes in the loading parameters Γ_high_, Γ_low_, t_high_ and t_low_, influence the structure of the maps?

### Discrete transitions underscore the molecular basis of junctional resilience

By coupling stochastic simulations (Figures 4a-f, blue) and analytical solutions (Figures 4g-1, red), we resolve how distinct regimes of cell-cell junction behaviour emerge from the molecular balance between linker association and dissociation under cyclic loading. In the rupture regime (Left column: Figure 4), linker dissociation dominates under sustained tension, leading to monotonic accumulation of bond ruptures and junction failure within a time shorter than thigh. In the full recovery regime (Right column: Figure 4), all broken bonds re-associate during each low-tension phase, fully restoring the equilibrium population of links and maintaining junction integrity over repeated cycles. In the slow damage accumulation regime (Middle column: Figure 4), the junction exhibits partial recovery between cycles, yet the cumulative net loss of linkers progressively shifts the junction towards failure. A phase space representation of the loss and gain of linkers can be found in Appendix 11, Figure 18.

**Figure 4:**
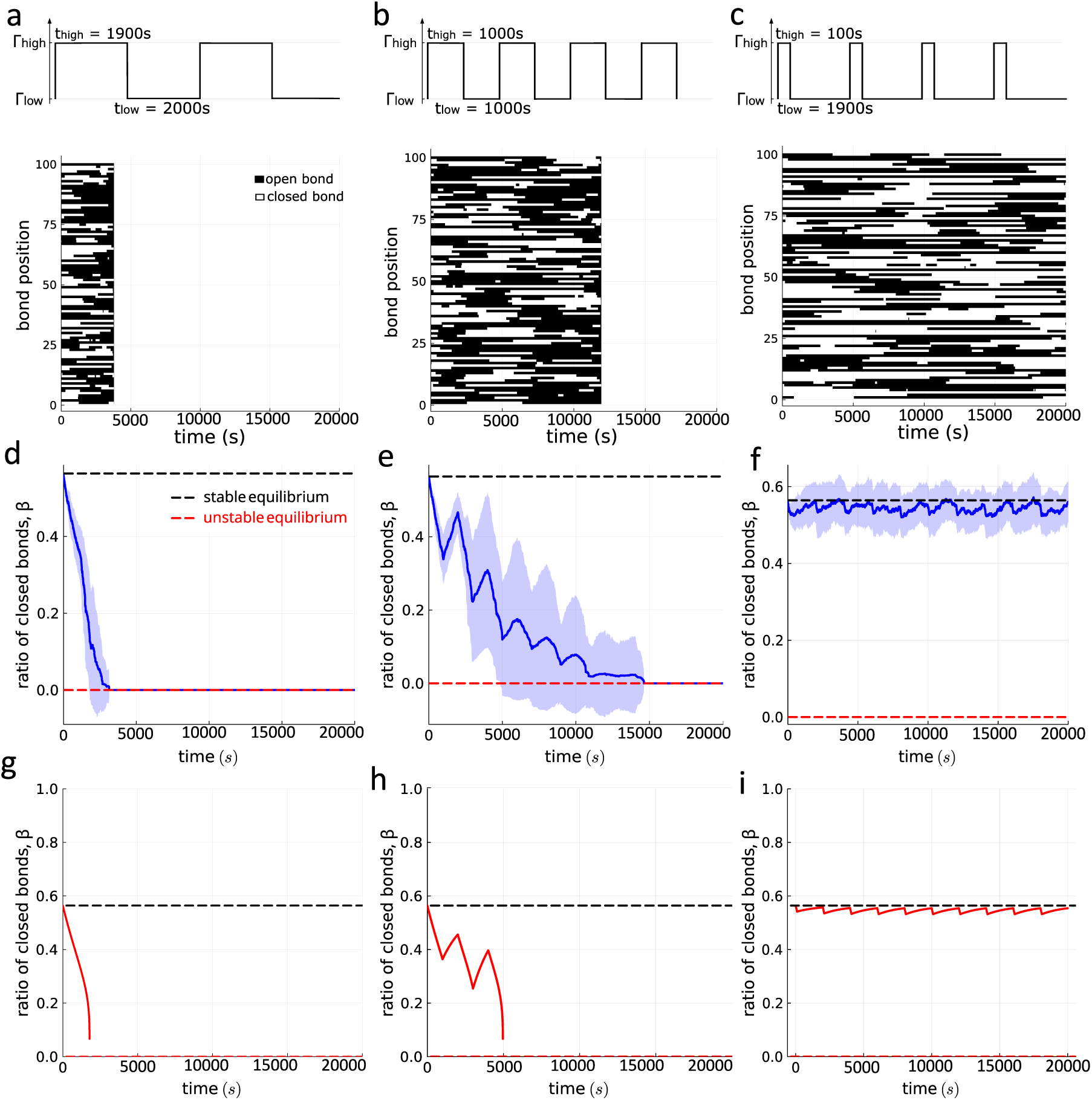
Balance of individual linker association and dissociation at the molecular scale predicts cell-cell junction response to cyclic loading. Left column: t_high_ > t^∗^, therefore all the bonds monotonically dissociate in the first cycle. Middle column: t_high_ ≈ t_low_, such that the relative ratios between the durations at high and low tension only allow for slow damage accumulation of the junction, resulting in delayed rupture. Right column: t_high_ ≪ t_low_, such that all the damage accumulated at high tension can be fully recovered and the bond ratio returns back to the equilibrium solution after each cycle. a-c) Top row: loading conditions for each of the three regimes: a: rupture, b: slow damage accumulation, c: recovery. Bottom row: Temporal maps of junction behaviour from stochastic simulations. A population of 100 links is shown. Black represents an open link and white a closed link, d-f) Temporal evolution of the ratio of closed bonds for each of the three regimes from stochastic simulations. Shaded regions denote the stochastic variability of bond dynamics across repeated simulations. The red dashed line is the unstable equilibrium and the black dashed line is the stable equilibrium g-i) Temporal evolution of the ratio of closed bonds for each of the three regimes from the analytical model.

Furthermore, there is a conserved pattern within the slow damage accumulation regime that can be distinguished by the presence distinct fringe-like bands in the analytical model. These bands mark discrete transitions in rupture time, corresponding to failure occurring after the first, second, third, or subsequent cycles (see Appendix 12, Figure 19). Mechanistically, this layered structure reflects the discrete nature of bond reformation. Each cycle at low tension, provides a quantized opportunity for linkers to re-associate, incrementally offsetting accumulated damage and thereby shifting the rupture event to a later cycle. As the number of cycles to failure increases, the spacing between successive fringes becomes progressively smaller and the junction approaches an asymptotic limit beyond which it transitions to the stable recovery regime. Although this structure is clearly expressed in the analytical model, it does not appear in the stochastic map, where fluctuations blur these discrete transitions. These results highlight that the observed macroscopic stability regimes are rooted in the emergent molecular-scale kinetics of individual linkers.

These molecular-scale observations show that the slow damage accumulation regime arises from the interplay between damage and healing within each loading cycle. To capture this balance quantitatively, we consider how the relative durations of the high- and low-tension phases compare with intrinsic timescales of rupture and repair. Specifically, the extent of damage accumulated per cycle can be estimated from the ratio of the applied high-tension duration, t_high_, to the rupture timescale, t^∗^, while the degree of recovery depends on the ratio of the low-tension duration, t_low_, to the repair timescale, t_e_. Additionally, the high- and low-tension amplitudes, Γ_high_, Γ_low_, can be normalised by the critical tension Γ_c_. Looking at the data through this lens removes any effects arising from the choice of specific loading parameters, allowing for comparisons across different loading conditions.

### Timescale normalization reveals a universal master stability map

Normalising the loading durations (t_high_ and t_low_) by the intrinsic rupture (t^∗^) and repair (t_e_) timescales reveals that the same three regimes, rupture, slow damage accumulation and healing, emerge across a wide range of absolute tension amplitudes (Figure 5). The general shapes of the boundaries between different regimes are qualitatively conserved across different combinations of loading parameters. When both Γ_high_ and Γ_low_ are far from Γ_c_, the system exhibits well-defined and robust regimes. This demonstrates that epithelial stability is governed by the relative interplay between intrinsic molecular kinetics and extrinsic loading timescales, rather than by the absolute magnitude of stress. For example, a tissue may persist longer under a high peak stress that is periodically relieved by low-tension intervals than under a continuous intermediate stress, highlighting that higher loads can be more sustainable when the loading pattern aligns with the intrinsic kinetic timescales. Despite differences in microscopic noise and individual bond variability, both the analytical and stochastic models produce equivalent regime boundaries when expressed in dimensionless timescales. This convergence indicates that the normalized map captures the fundamental physics of rupture–repair competition, independent of model implementation. Moreover, the slope of the initial linear region between healing and slow damage accumulation was found to be robust to changes in tension. When perturbing the dynamics of linker kinetics by varying k_on_ and k_off,0_ while preserving their ratio, the structure of the normalised stability maps also remained robust (see Appendix 10, Figure 17). The model dynamics are determined primarily by the relative kinetics of bond association and dissociation, rather than the absolute rate constants. This highlights a key strength of the framework: by normalizing mechanical inputs by intrinsic molecular timescales, all loading conditions collapse onto a single master stability map.

**Figure 5:**
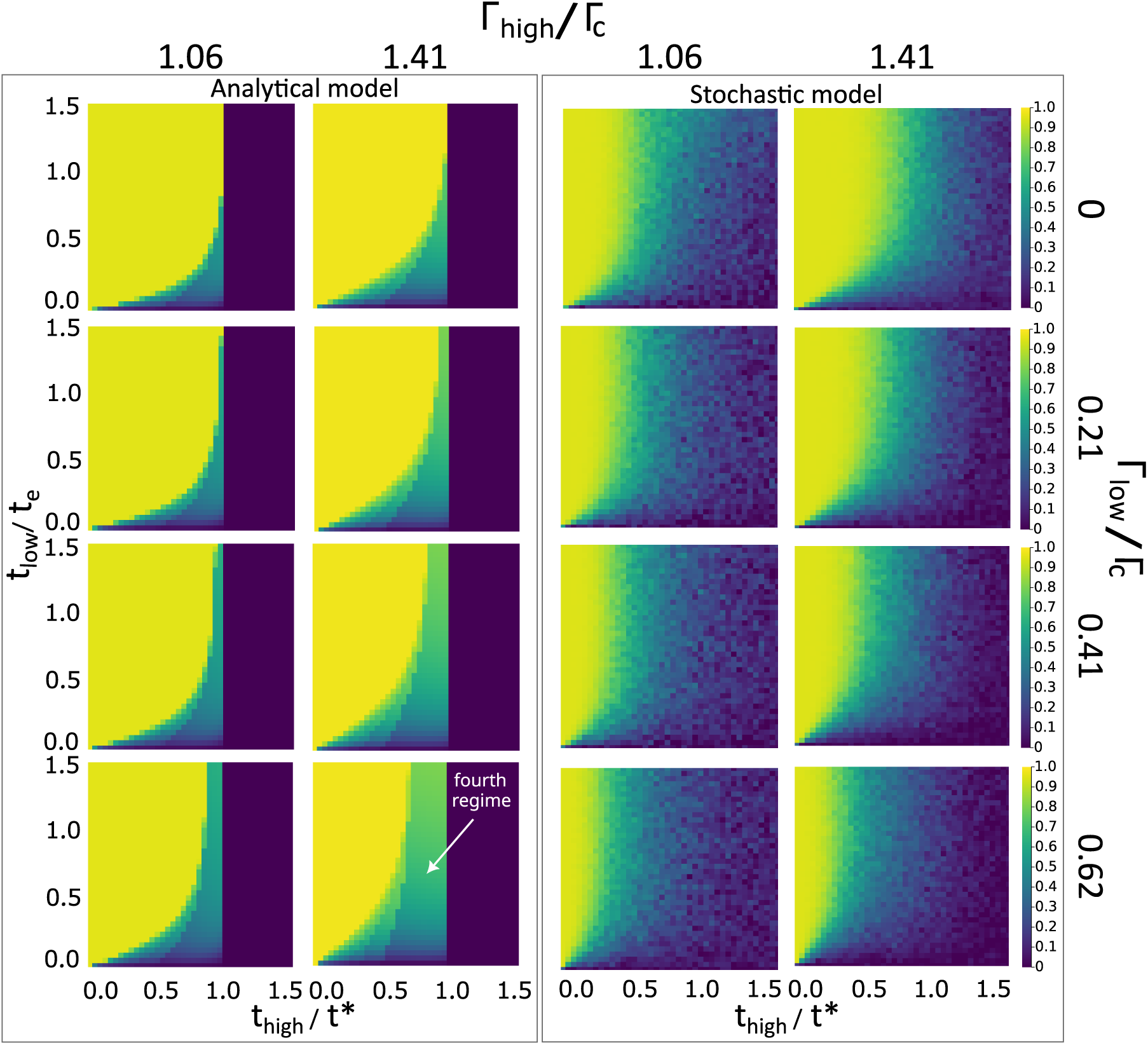
Distinct regimes of epithelial stability, with conserved characteristics. Different combinations of Γ_*high*_ ranging from 0.09*Nm*^−1^ to 0.12*Nm*^−1^ and Γ_*low*_ ranging from 0*Nm*^−1^ to 0.0525*Nm*^−1^. For each combination, the stability graph is computed for different ranges of tlow and thigh. In these graphs, *t*_*low*_ is normalised by the repair timescale, *t*_*e*_, and *t*_*high*_ is normalised by the rupture timescale, *t*∗, of the corresponding tension values. Columns one and two capture the dynamics of the analytical model and columns three and four capture the dynamics of the stochastic model. The white arrow points to the fourth regime that emerges for non-zero Γ_*low*_ values.

Another key observation is that as either value approaches Γ_c_, the phase boundaries become less distinct and more sensitive to stochastic fluctuations. Notably, the closer Γ_low_ is to Γ_c_, the broader and more diffuse the transition between the rupture, slow damage, and recovery regimes becomes. We find that increasing Γ_low_ while keeping Γ_high_ constant, progressively reduces the size of the healing region. In this case, the effective window for rebinding decreases, especially as the repair timescale t_e_ becomes much longer relative to t_low_. Consequently, junctions spend a larger fraction of the cycle under damaging conditions, shifting the balance from reversible to irreversible failure. Interestingly, when Γ_low_ is set to non-zero values, we identify a fourth stability regime (white arrow, Figure 5), which appears as a rectangular band within the slow damage accumulation regime and extends into the healing phase space for increasing values of Γ_low_. In this regime, the linker loss during the first high-tension phase drives the system across the unstable fixed point, placing it within the diverging regime of the model. In this case, even when the tension is reduced to Γ_low_, the junction continues to accumulate damage, albeit at a slower rate. In this fourth regime, initial conditions become especially important, as small differences in the initial bond population can determine whether the junction crosses the unstable solution or not (see Appendix 9, Figure 16).

The normalized framework enables direct comparison across tissues or conditions by mapping experimental measurements (e.g., cycle frequency and duty ratio) onto the same dimensionless axes. Tissue stability can therefore be predicted independently of specific parameter choices, making the framework broadly applicable to biological systems with diverse molecular dynamics and different loading conditions.

## DISCUSSION

Our experiments showed that when epithelial monolayers are held under a sufficiently high constant tension, they progressively deform and ultimately rupture, with higher loads producing faster failure and smaller strains at the rupture point. We then investigated the reversibility of damage by subjecting the monolayer to cycles of controlled supercritical and subcritical tension. Under repeated cycles of high and low tension, the monolayers’ lifetime at high load and overall deformation increased markedly, compared to constant loading. To determine whether this enhanced resilience arises from emergent physical behaviour rather than active adaptation, we turned to a simple micromechanical model of a linker population with slip-bond kinetics. Under cyclic loading, the model predicted a rich range of behaviours, ranging from fast rupture to stable adhesion. When a junction approached rupture, damage accumulated continuously as intercellular links weaken and separate. Periods of low tension appeared to allow partial recovery of links, enabling the tissue to withstand high stresses for longer. A distinct regime of slow damage accumulation accounts in particular for our experimental observation of delayed rupture under cyclic loading. The key to understanding these dynamics lies in quantifying two timescales: the rupture timescale at high load, corresponding to the time required for all the bonds in a population to dissociate, and the repair timescale at low or zero load, over which the bond population returns to a kinetic equilibrium. The magnitudes of these timescales, relative to the durations of loading and rest, control the outcome of exposure to cyclic loading.

Our results reveal emergent damage accumulation and healing behaviours arising from the coupled dynamics of cellular mechanics, tissue rheology and junctional remodelling. In response to deformation, the rheology of the tissue sets the stress magnitude to which cell-cell junctions are subjected. Although the intercellular adhesion complexes (cadherin-catenin, desmosome-desmosomal cadherins) linking the cytoskeleton of adjacent cells contribute negligibly to bulk deformation, the dynamic remodelling of the intercellular linker population under force plays a critical role in setting tissue strength and controlling the onset of failure. This coupling highlights that studying the mechanical behaviour of epithelial tissues requires a holistic approach, encompassing the entire force transmission pathway, from the molecular linkers that mediate adhesion to the collective deformation of all the cells in the tissue. The stochastic turnover of adhesive bonds shapes the transition between reversible and catastrophic damage accumulation. Importantly, the emergent rupture and repair behaviour can arise naturally from the binding and unbinding kinetics of molecular linkers under load. Our results also highlight the broader significance of cellularisation. Tissues’ organisation into discrete mechanical units enables the independent tuning of strength and rheological properties, providing an advantage that cannot be captured by a single continuum property, but emerges from collective molecular interactions at cell-cell interfaces (26).

When tissues operate near the boundaries separating rapid rupture, effective repair, and slow damage accumulation, their response becomes highly sensitive to repair kinetics, making the repair timescale a key determinant of tissue integrity. This repair timescale is distinct from the cellular processes involved in wound healing, such as lamellipodial protrusion or purse string closure (27). Our findings outline a strategy to probe tissues’ dynamic response under cyclic loading, providing a means to address how epithelial tissues preserve their integrity under repeated mechanical perturbations. Systematic variation of Γ_high_, Γ_low_, t_high_ and t_low_ enables direct measurement of rupture onset and the gradual accumulation of damage over successive cycles. However, direct experimental characterisation of the repair timescale remains challenging, as it involves molecular processes occurring below optical resolution and over a wide range of timescales (28). In principle, the repair timescale, t_e_, can be estimated indirectly by taking advantage of the broadly conserved normalised stability maps. The experimentally measured t^∗^ and applied t_high_ and t_low_ can be used to locate the loading protocol on the map, from which t_e_ can be inferred relative to the low-tension phase. In the case of our MDCK monolayer protocol, the minute-scale t_low_ confined the tissue to a partial-repair regime. This observation suggests that the repair timescale *t*_*e*_ in MDCK monolayers is likely on the order of hours. While direct measurements of junctional repair timescales remain scarce, prior studies of E-cadherin and associated p120-catenin complexes show turnover and membrane re-incorporation occurring on the timescales of tens of minutes (12, 29, 30). Nevertheless, we anticipate that advances in single-molecule imaging and protein-protein interaction may enable direct measurement of the open or closed configuration and lifetime of individual adhesion proteins under controlled mechanical load (28, 31). Such measurements would allow quantitative validation of the predicted coupling between linker kinetics and tissue-scale forces.

Our analysis focuses on a single population of adhesive linkers with uniform kinetics, providing a simple approach to connect molecular dynamics to macroscopic mechanical behaviour. In reality, healthy tissues draw on a broad range of adhesion proteins (cadherins, desmosomal cadherins, tight junction proteins, JAMs, etc) with different strengths and dynamic properties, enabling junctions to form and stabilize even under minimal or oscillatory loads (32). Such molecular diversity may provide a basis for resilience, allowing different intercellular proteins to contribute to cell adhesion (33). Incorporating heterogeneity in the kinetics of individual linkers in the framework, offers a testbed for this hypothesis while paving the way towards a multiscale description that captures the added complexity and reveals how molecular architecture shapes tissue strength.

The recovery response we highlight is relevant beyond biological tissues, offering direct analogies to a broad class of soft materials such as self-healing polymers and dynamic hydrogels (34–37). Although these materials have different structural components, they rely on comparable reversible binding mechanisms. In these systems, the introduction of reversible linkers, such as dynamic covalent bonds, hydrogen bonds and metal-ligand complexes, enables materials to dissipate energy and recover after damage (34). Our model captures the same conceptual principle, showing that macroscopic resilience emerges from the collective dynamics of microscopic reversible bonds. The molecular dynamics seen in epithelial junctions could guide bioinspired engineering of synthetic networks with tunable strength and reversible adhesion (38). For instance, tuning linkers’ force sensitivities or timescales to mimic biological adhesion could provide both rapid healing and long-term durability. Similar design strategies are now being explored in self-healing polymer networks and hybrid hydrogels, where hierarchical or multi-link architectures enhance performance under cyclic loading (35, 36). Ultimately, the physics of epithelial cohesion offers a foundation for engineering synthetic materials that are simultaneously strong and self-repairing.

The cyclic loading periods used in this study, on the order of 1000s on and 1000s off, correspond to those of processes such as intestinal peristalsis or bladder filling and voiding, which occur over tens of minutes and involve substantial mechanical stress (5). Physiologically, our experiments and simulations indicate that soft biological tissues can tolerate extended periods of overload without rupture, provided these are interspersed with phases of low tension that permit recovery. Locating which regime tissues operate in and whether they are close to boundaries may therefore help assess their susceptibility to mechanical failure and disease (8, 39–42). Tissues positioned close to regime boundaries may be particularly vulnerable, as modest random fluctuations in tension or junctional stability can tip the system into failure. Such a mechanical vulnerability could contribute to multifactorial diseases, such as Crohn’s disease and epidermolysis bullosa (43, 44). Comparison of the predictions of our simulations to experimental data may highlight the presence of active cellular responses to damage. Indeed, enhanced recovery, adaptive adhesion, or delayed failure, likely reflect the activation of active feedback and complex signaling pathways that modulate adhesion in response to load. Therefore, the subtle balance between damage accumulation and healing may be physiologically relevant and important for assessing the health of tissues.

## CONCLUSION

Overall, contrary to the notion of stable junctions conveyed by their continued presence in timelapse videos of epithelia, the intercellular bonds that underlie these junctions are inherently dynamic, with the number of closed intercellular bonds fluctuating as a function of applied load and time. The interplay between applied force cycles and intrinsic molecular kinetics governs the transition between rupture and repair. These insights pave the way for a more predictive understanding of epithelial tissue mechanics that incorporates both loading dynamics and molecular response timescales.

## MATERIALS AND METHODS

### 1 Experimental *in vitro* model

#### 1.1 Cell culture

Madin-Darby Canine Kidney (MDCK) wild type cells were cultured at 37^◦^C in an atmosphere of 5% CO2 in DMEM (1X) (Thermo Fisher) supplemented with 10% fetal bovine serum (FBS, Sigma-Aldrich), 2.5% of 1M HEPES buffer (Sigma-Aldrich) and 1% penicillin-streptomycin (Thermo Fisher). Cells were passaged at 1:8 ratio every 4 days using standard cell culture protocols and disposed of after 15 passages. Mechanical experiments and imaging were performed in Leibovitz’s L15 (Thermo Fisher) supplemented with 10% fetal bovine serum (FBS, Sigma-Aldrich) and 1% penicillin streptomycin (Thermo Fisher).

#### 1.2 Stretching devices

The stretching devices were microfabricated following the protocol described in (21), with some adaptations. The materials required to make each device are: borosilicate glass, nitinol wire (two thicknesses), tygon rings (two thicknesses) and UV glue. Each assembled device consists of one stiff rod covered by a glass capillary and one compliant soft rod. Two rings of tygon of different thicknesses cover the extremities of the two wires, such that epithelial cell monolayers can grow between the test rods. The step-wise assembly involves (i) local melting of the borosilicate glass capillaries into a U-shape using a micro-pen blow torch (Inoda PT-200 Pro torch), (ii) using pliers to cut the bent capillary into one U-shape (5mm) and one straight section (25mm), cutting the thick wire (30mm), and the thin wire (30mm), using scissors to cut the thick tygon (5mm) and the thin tygon (5mm), (iii) connecting the parts of the devices and securing in place using UV curing glue, (iv) curing for 30min. Each device was secured to the bottom of a 50mm petri dish by applying biocompatible adhesive glue on the U-shape glass base, minimise any lateral movement during the experiment. During the gluing step, a short glass capillary was placed beneath the tygon tips to slightly elevate the device, preventing contact with the bottom of the dish.

#### 1.3 Preparation of suspended MDCK monolayers

The protocol for generating suspended MDCK monolayers is based on the work of Harris 2013 and Duque 2024 (10, 21). Cells were cultured on a sacrificial collagen substrate. Collagen type 1A was reconstituted on ice, by mixing 50% collagen (Cellmatrix type I-A, Nitta Gelatin), 20% sterile milli-q, 20% of 5X DMEM and 10% reconstitution buffer 50 mM NaOH solution in sterile water, 200 mM HEPES and 262 mM of NaHCO3. A 10uL droplet was placed between the device’s test rods. The devices were then placed in an incubator for drying over 60 - 75 minutes at 37^◦^C. This was followed by a rehydration step with 10*μ* l of culture medium for 30 minutes at 37^◦^C. Approximately 40,000 cells were seeded on the rehydrated collagen and incubated at 37^◦^C and 5%CO2 for 30 - 45min. Lastly, 55ml of culture medium was added to completely submerge the test rods and the monolayers were left in the incubator to grow for approximately 72h, or until the monolayer extended from one rod to the other, covering the collagen substrate as well as each rod partially. Before mechanical testing, the collagen was digested enzymatically, using collagenase type-II (Worthington Biochemical) with imaging medium to reach a final concentration of 250 units per mL. The culture medium was exchanged for the collagenase solution and was placed at 37^◦^C for 1h. Before testing, the collagenase solution was exchanged for the imaging medium. Under these conditions, the cells remained healthy and retained their characteristic epithelial apico-basal polarity for 3h.

#### 1.4 Force measurement

A petri dish containing a stretching device and suspended monolayer was placed on the stage of an inverted brightfield microscope (Olympus IX71). The stiff rod was attached to a motorized micromanipulator (M126-DG1) controlled through a C-863 controller (Physik Instrumente) via a LabVIEW code (National Instruments). During experiments, force was applied to the tissue by stretching it through controlled movements of the micromanipulator (10, 12, 21). The soft rod was attached to the tip of a force transducer (SI-KG7A, World Precision Instruments) which deflects relative to the deflection of the soft wire. The deflection is converted into a digital signal and recorded as voltage data (USB-1608G, Measurement Computing). Images were taken every 6s using a x2 objective (PLN2X, Olympus) and a camera (Point Grey Grasshopper3 U3, GS3-U3-60QS6M-C). The applied force can be measured by monitoring the bending of the soft test rod, detected by the force transducer tip. At the end of each experiment, the monolayers were broken and the rod’s position after failure was used to define the zero-force geometry. Increments of known deformation were applied manually to obtain a set of six voltage-deformation values. The voltage-to-stress conversion was done using a standardised calibration procedure from Duque 2024 (10) (see Appendix 1).

#### 1.5 Stress controller

Constant stress was applied by tuning the position of the motorized micromanipulator, using a Proportional-Integral-Derivative (PID) controller (Figure 6). Before each creep experiment, a pre-calibration was performed on each device, to obtain a voltage-displacement relationship and a unique conversion factor, *α*. The target voltage was computed by taking the difference between the target stress and prestress divided by the calibration coefficient. The error signal, E(t) was computed by subtracting the measured voltage from the target voltage and is fed into the PID controller. The controller minimises this signal by adjusting the strain applied to the sample via the motorized micromanipulator. The PID controller adjusts the micromanipultor’s position at a maximum rate of 0.1mm/s which is equivalent to a maximum strain rate of 10%s^−1^. This rate cap was chosen to minimise the effects of strain-rate-dependent rupture that have been reported in the literature (10). The controller parameters, K_P_, K_I_, K_D_, were empirically tuned for stable convergence to the target stress. The optimised controller parameters are found in table 1. To assess the controller’s performance, a post-experiment calibration was performed following each test. Callibration coefficients before and after the experiment were compared by plotting them against each other. The resulting data closely aligned with the identity line which fell within the confidence interval of the data, indicating that the pre-experiment calibration coefficients were reliable for computing the target voltage (see Appendix, Figure 7).

**Figure 6:**
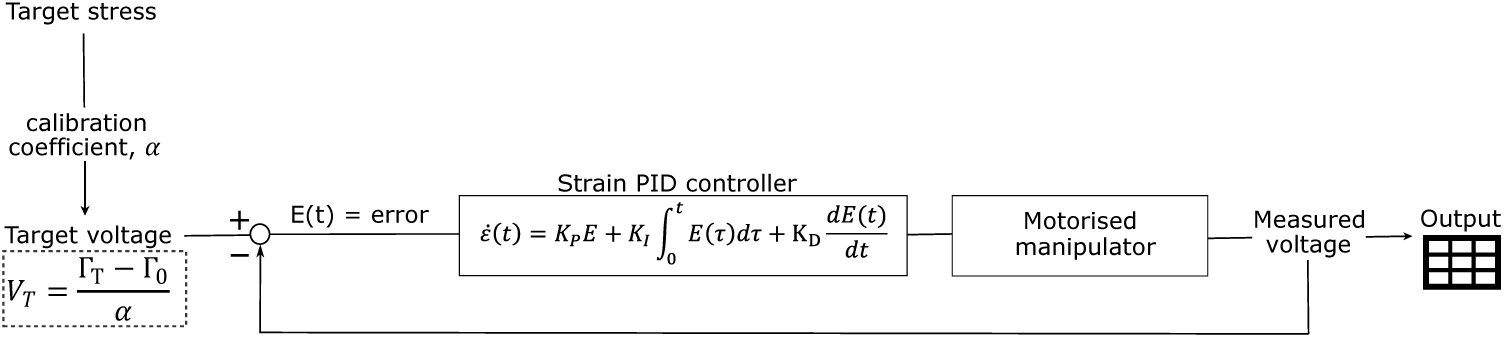
PID controller logic. The target voltage is calculated using the target stress, Γ_T_, the prestress, Γ_0_ and the pre-calibration coefficient, *α*. The error signal, given by subtracting the measured voltage from the target voltage, is then minimised through the PID controller which adjust the strain by adjusting the motorised manipulator position. At each timestep the data output is recorded on a text file.

**Table 1:** Stress controller parameters.

| Parameter | Value |
| --- | --- |
| $K_P$ | 1.2E-1 |
| $K_I (1/s)$ | 0 |
| $K_D (s)$ | 5E-5 |
| control velocity (mm/s) | 1E-1 |

### 2 Analytical model

The analytical slip-bond model is based on Bell’s theoretical framework of cell adhesion (14). The governing analytical equation 2 is solved in Julia by creating an ordinary differential equation problem, using the Tsit5() Runge–Kutta equation solver from the OrdinaryDiffEq package (45). The relative and absolute tolerances are set to 1*e*^−8^ and the step, dt, is set to 1*e*^−2^. The rupture timescale, t∗ was initialised at the stable bond ratio for zero tension (Equation 4), where k_on_ is the association constant rate and k_off,0_ is the dissociation rate at the start of the simulation. An alternative initialisation considered was starting at 100% bonds closed, however this is not a physiologically representative condition, as there is continuous turnover of intercellular junction proteins and would likely lead to an overestimation of the rupture timescale. The repair timescale, t_e_, was initialised at the critical ratio of bonds, which corresponds to the ratio of bonds at critical tension, Γ_c_ and was computed by calculating the characteristic time of the exponential. For the cyclic loading condition, the model was also initialised at the critical ratio of bonds.

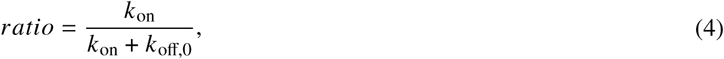

The slip bond model parameters (table 2) were obtained by fitting the model’s fracture timescales to experimental fracture timescales using the root mean squared error (RMSE) in tension. Optimisation was performed using the NelderMead algorithm in Julia’s Optim package (46).

**Table 2:** Slip bond model parameters.

| Parameter | Value |
| --- | --- |
| $k_{on}(1/s)$ | 3.47E-4 |
| $k_{off,0}(1/s)$ | 2.68E-4 |
| $f_0(N/m)$ | 2.49E-1 |

### 3 Stochastic model

The stochastic model is based on the same governing equations as the analytical model, applied to a population of links with slip-bond properties. A junction is defined with N=100 links and each link can independently associate or dissociate with probabilities defined by equations 1a and 1b. The junction dynamics are simulated using a Monte Carlo scheme, initialised with a random distribution of link states evolving in discrete probability steps. Each run results in either complete rupture of the junction (all links dissociate) or in reaching the maximum duration (three times longer than the largest repair timescale), in which case the junction is classified as healed. All stochastic simulations were perfomed using the open-source Julia package CellAdhesion.jl (47). Initialisation conditions were set by assigning bond states randomly across a network of 100 links, each representing an individual molecular connection. For each condition (rupture, healing or cyclic loading) the proportion of initially closed bonds was determined based on the corresponding ratio used in the analytical model. In each case, the number of closed bonds at the start of the simulation was calculated by multiplying the total number of links by the corresponding ratio, with the positions assigned randomly across the network to avoid spatial bias.

### 4 Phase map generation

Phase maps were generated for different combinations of Γ_high_ and Γ_low_ each with 40×40 *t*_high_ x *t*_low_ combinations. The range of 40 possible *t*_high_ values was scaled from 0.01 to 4.5 times the rupture timescale. Similarly, the range of 40 possible *t*_low_ values was scaled from 0.01 to 1.5 times the repair timescale. For the stochastic model phase maps, the same coordinates were selected, but 100 model simulations were done at each coordinate and the mean was plotted. The maximum simulation duration was set to 10 times the longest loading cycle period, corresponding to 10 times the sum of one and a half times t_high_ and one and a half times t_low_. This design choice ensured that any events classified as healing were sufficiently different from rupture or slow damage accumulation regimes. The simulation timestep, dt, was set to one-hindredth of the smaller of t_high_ and t_low_, providing adequate temporal resolution across all tested regimes.

## AUTHOR CONTRIBUTIONS

All authors conceived the project and wrote the paper. E.P. performed the experiments, imaging and analysis. G.C. advised on the analysis of experimental data. E.P. implemented the analytical and stochastic modelling. A.B. and A.J.K. designed the stochastic model and advised on the analysis of computational data. A.B., G.C. and A.J.K. oversaw the entire project. All authors discussed the results and the paper.

## ACKNOWLEDGMENTS

We thank the past and present members of the Charras and Kabla laboratories for discussions. We wish to acknowledge Nargess Khalilgharibi and Jonathan Fouchard for technical support with LabVIEW development and Lucia Baldauf for her important contribution in laboratory training and insightful discussions. E.P. was supported by the Engineering and Physical Sciences Research Council Centre for Doctoral Training in Sensor Technologies for a Healthy and Sustainable Future [EP/S023046/1], G.C. was supported by an sLoLa grant from the British Biotechnology and Biological Sciences Research council (BBSRC, grant no. BB/V019015/1). A.B. was supported by the European Research Executive Agency via the MSCA Postdoctoral Fellowships (BIOFRAC, Project number: 101105376).

## APPENDIX

## 1 Force measurement: calibration procedure

Force can be calculated using the cantilever beam equation (5), where k is the rod’s stiffness (which depends on the material and geometrical properties of the wire) and dw is the deflection. The value of k can be computed using equation 6, using the Young’s modulus, E, the second moment of area, I, and the length, L, of the wire.

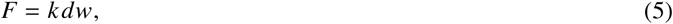

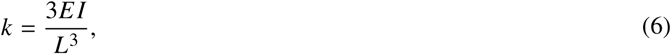

Using six pairs of voltage and deflection (V, dw) a linear fit is used to determine the conversion factor from Volts to Newtons, given by equation 7.

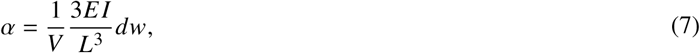

Using a 2D thin sheet approximation and equation 8, the force of each monolayer was normalised by the average width, w0, to calculate tissue tension. The thickness of the monolayer is ∼ 10*μ*m which is two orders of magnitude smaller than the length and width of the monolayer, justifying our approximation. The cross-sectional area was computed by multiplying the initial length by the average width. The average width was found by averaging the width at each of the arms of the device and in the middle of the monolayer.

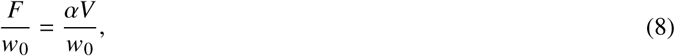

Finally, the monolayer’s pre-tension was calculated using two bright-field microscopy images, one from the start and one from the end of the experiment. A 250×150p×2 area was cropped around the flexible rod to measure its displacement Δx that can be used to find the pre-tension, using equation 9. The actual tension values were computed by subtracting the pre-tension offset.

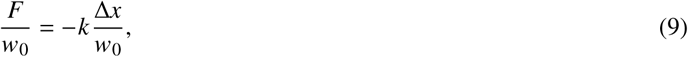

**Figure 7:**
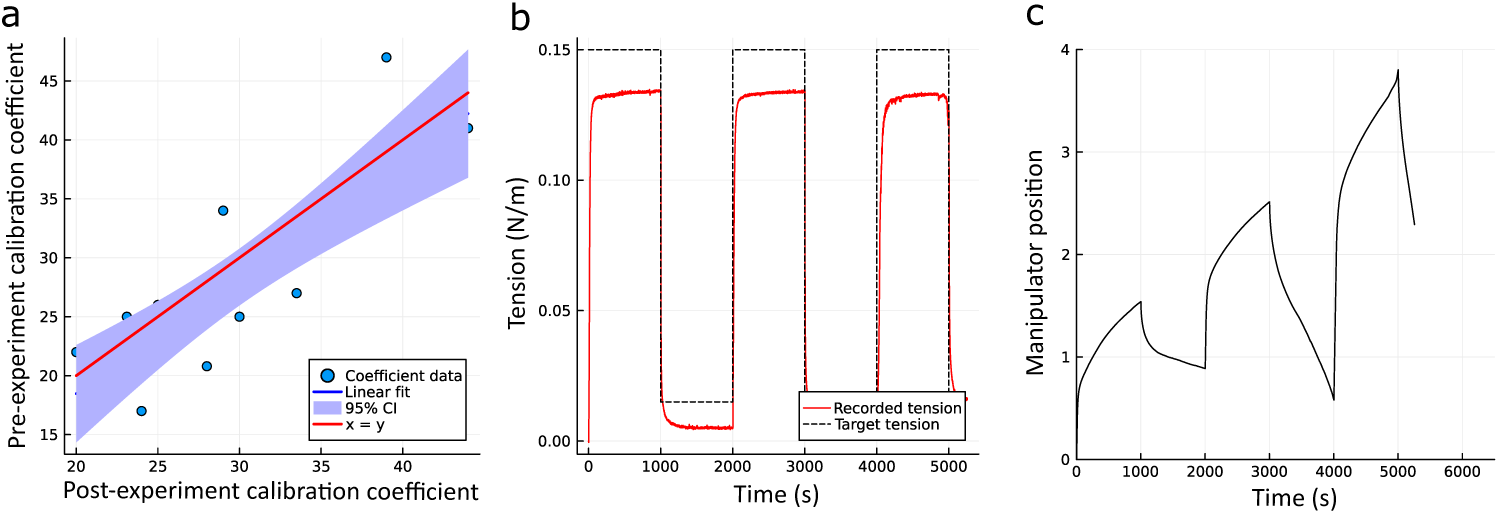
PID controller performance. a) Comparison of pre- vs post-experiment calibration coefficients. Data points fall close to the identity line (x=y, red), and the linear fit lies within the 95% confidence interval (shaded, blue), indicating that pre-experiment coefficients reliably estimate post-experiment values. b) Target tension (black, dashed) compared to recorded tension (red, solid) during cyclic loading, showing that the controller accurately tracks the target setpoints across cycles. c) Manipulator position over time, illustrating dynamics adjustment of the deformation in each cycle to achieve the target tension, demonstrating active feedback control during repeated loading.

**Figure 8:**
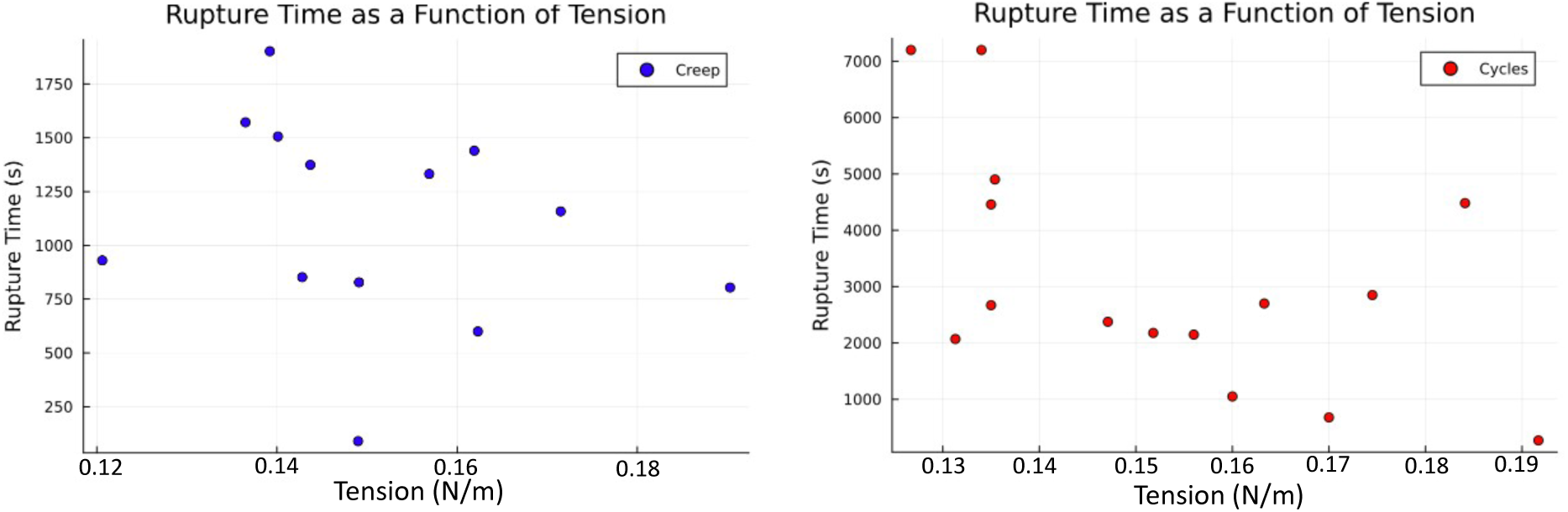
Rupture time as a function of applied tension (Γ_high_) for creep (left) and cyclic loading (right) tests. For creep, rupture times show no clear monotonic dependance across the examined tension range (0.12 - 0.195N/m). In the cyclic loading case, a slight decreasing trend with increasing tension is observed, though variability remains large. These observations support including tests spanning this full tension range.

## 2 Distribution of experimental and model fracture times for target tension

## 3 Strain response to creep under cyclic tension

**Figure 9:**
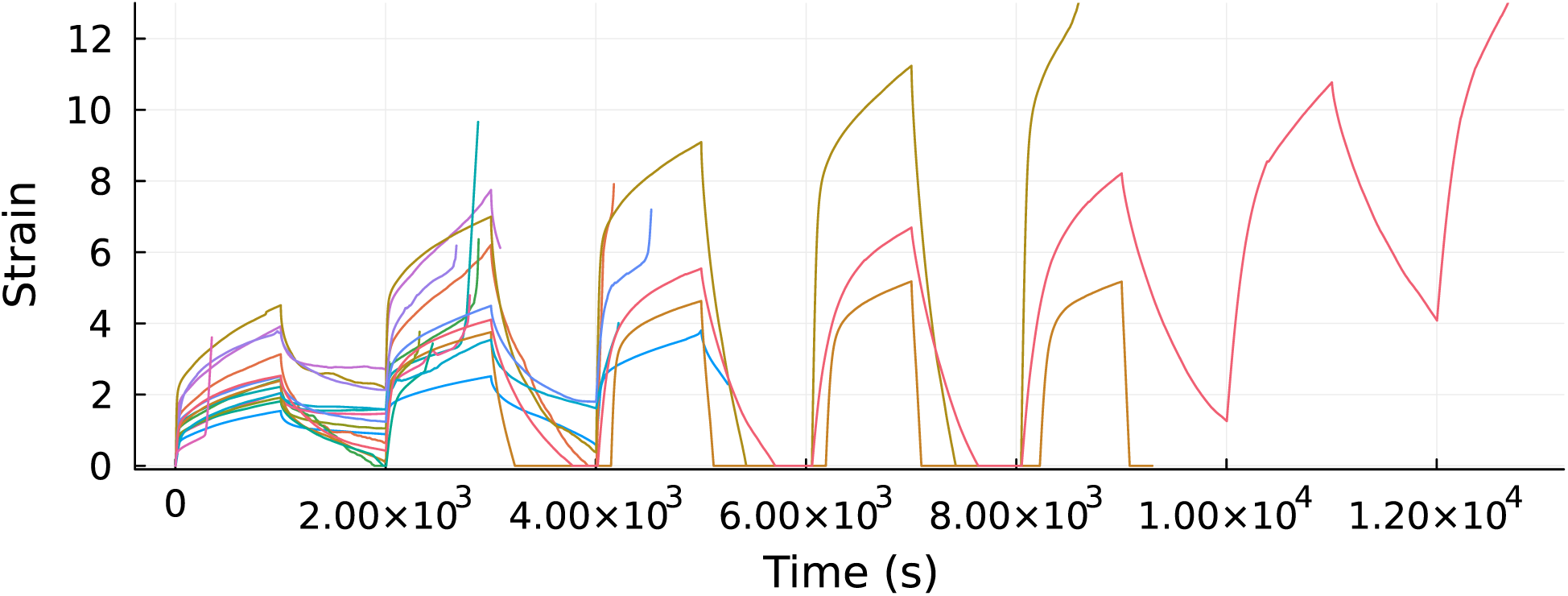
Experimental strain response during cyclic creep loading. Each curve represents an individual loading–unloading cycle measured under constant applied stress. Later cycles diverge markedly, indicating the onset of irreversible deformation and rupture within the epithelial monolayers.

## 4 Slip bond parameters: fitting and optimisation details

Fracture times were not used directly, as they diverge near the critical tension and would yield undefined RMSE values. For each experimentally measured fracture time, the analytical model was used to find the tension value producing the same rupture timescale. The squared difference between this predicted tension and the experimental value was computed, and the square root of the mean of these squared differences gave the first RMSE value. To enforce accuracy at the main experimental condition, we identified the mean fracture time corresponding to 0.145Nm^−1^, used the analytical model to predict the tension yielding the same fracture time and added the squared deviation between this predicted tension value and 0.145Nm^−1^ as a second error term. The combined objective function was defined using equation 10, were the factor of three balances accurate reproduction at 0.145Nm^−1^, with fidelity across the full fracture-time profile.

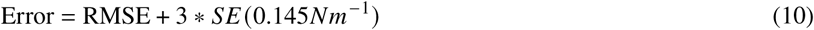

The analytical model was evaluated for 100 logarithmically spaced tension values between 10^−5^ and 10N/m, capturing rupture timescales from near zero to 4000s. This range ensured stable optimisation and uniform sampling of the model’s exponential decay behaviour. Multiple initialisations were tested and the parameter set yielding the lowest error was selected. Initial values for k_on_ and k_off_ were 3*e* − 3, 3*e* − 4, 3*e* − 5 and for f_0_ inital values were 0.0055N/m, 0.055N/m, 0.55N/m. These ranges include parameters previously obtained from epithelial monolayer strain-ramp loading experiments (10) and cover three orders of magnitude. Parameters were optimised in logarithmic form to enforce positivity without explicit bounds.

**Figure 10:**
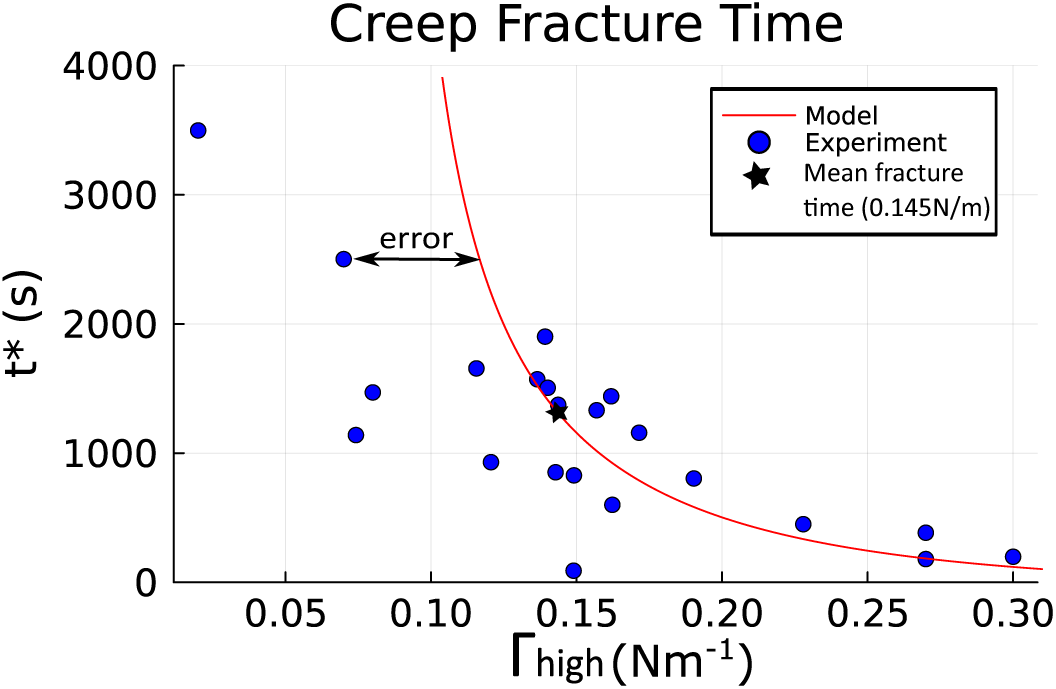
Creep fracture time as a function of applied tension. Blue points show experimental data, the red curve shows the model, and the black star marks the mean fracture time at 0.145Nm^−1^. The horizontal arrow illustrates the definition of error in tension, which forms the basis of the RMSE calculation. Overall model error is computed as RMSE plus a weighted term ensuring accurate reproduction of the 0.145Nm^−1^ condition.

## 5 Slip-bond model parameter fit to experimental data

**Figure 11:**
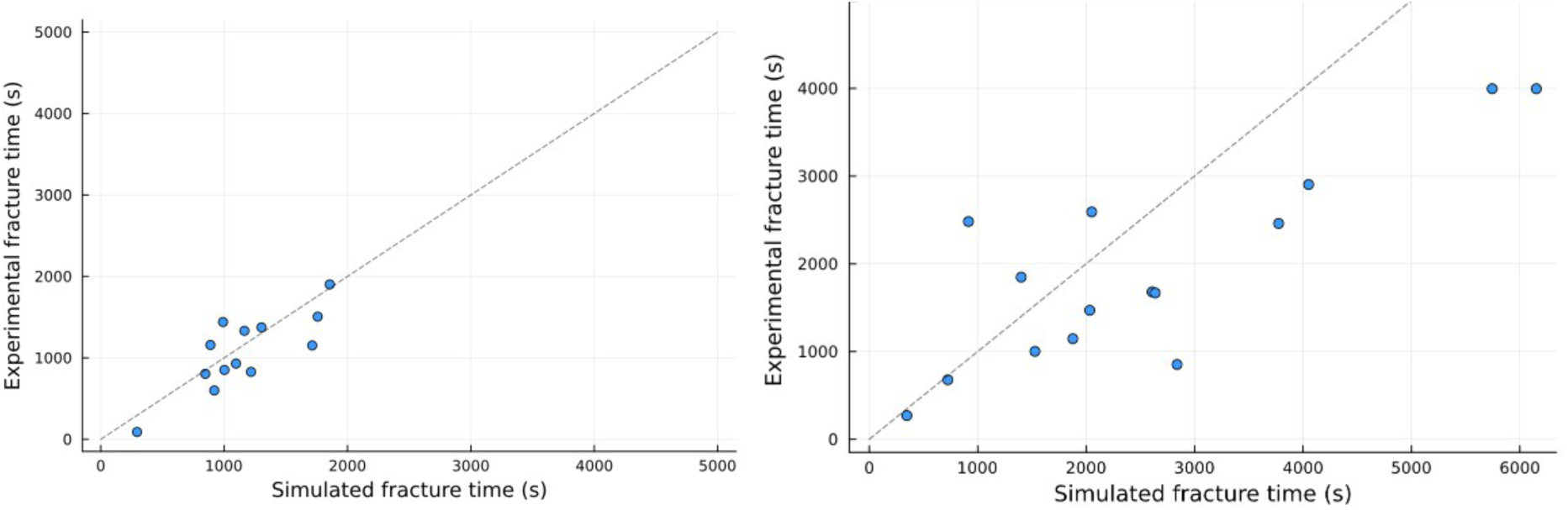
Comparison of simulated (stochastic model) and experimental rupture times for creep (left) an cyclic loading (right). Each point represents an individual experiment, with the dashed line indicating perfect agreement between simulation and experiment.

## 6 Simulated rupture and repair timescales as a function of tension for different bond numbers

**Figure 12:**
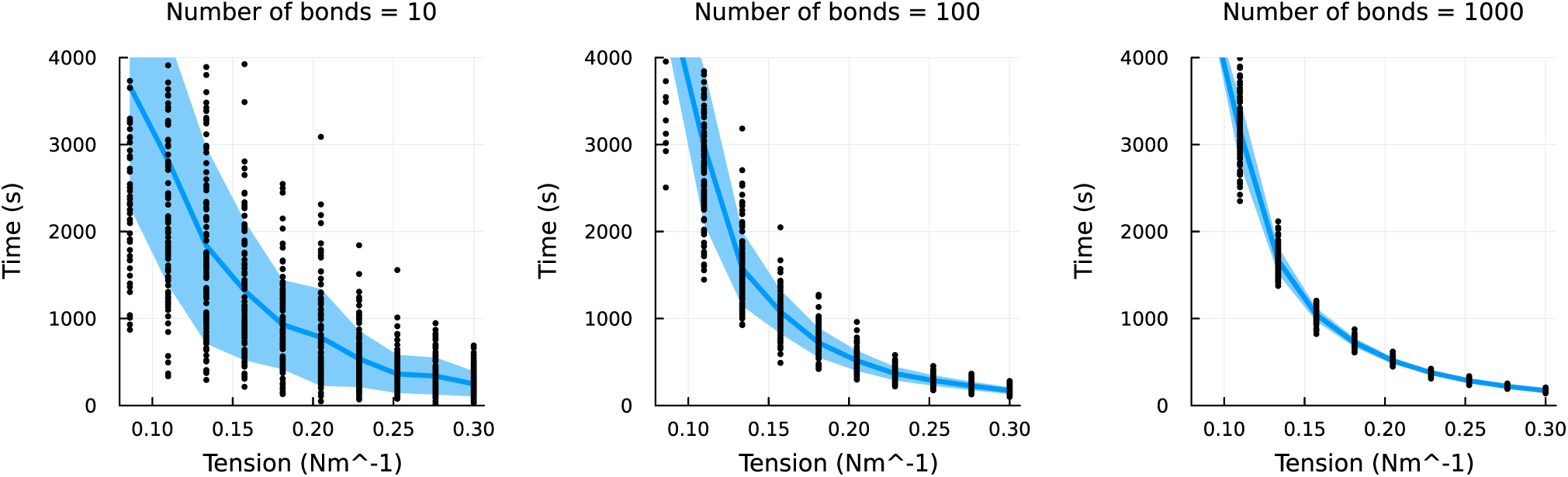
Rupture time as a function of applied tension for 10, 100, and 1000 bonds. Each black point represents an individual stochastic simulation, and the shaded blue regions indicate the standard deviation across runs. Increasing the number of bonds decreases the variability. N=100 bonds matches the noise of previous experimental data (10).

**Figure 13:**
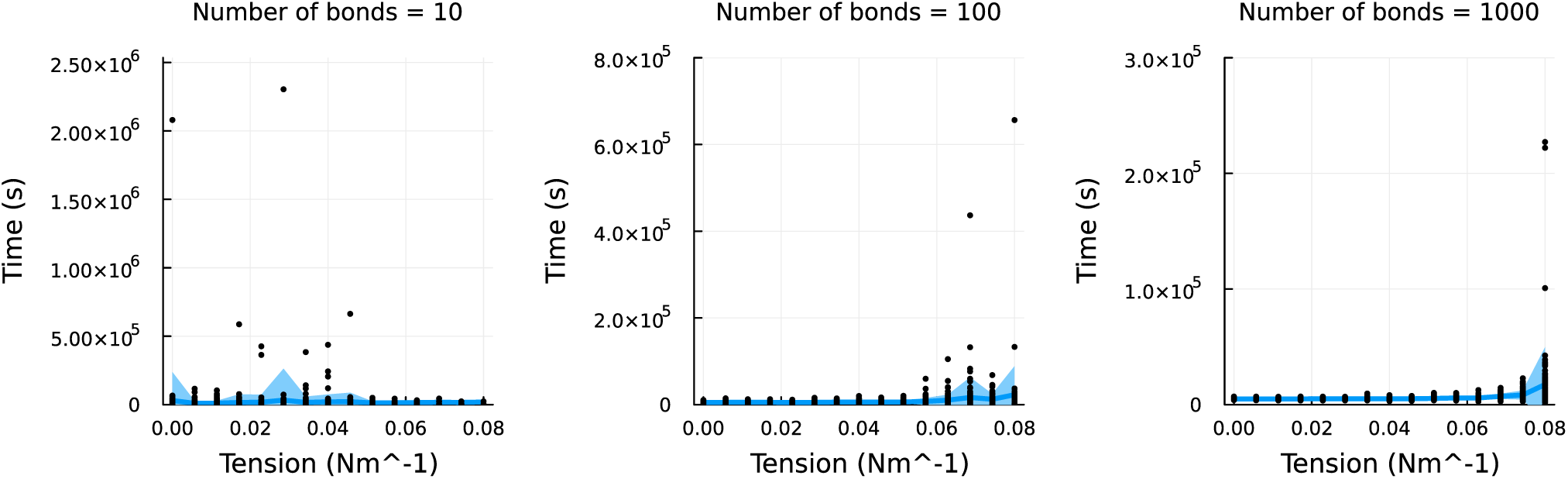
Recovery time as a function of applied tension for 10, 100, and 1000 bonds. Each black point represents an individual stochastic simulation, and the shaded blue regions indicate the standard deviation across runs. Repair is more variable with fewer bonds with certain points exceeding tenfold slower recovery times for N=10. Increasing the number of bonds decreases the variability. N=100 bonds matches the noise of previous experimental data (10).

## 7 Recovery dynamics close toΓ_*c*_

**Figure 14:**
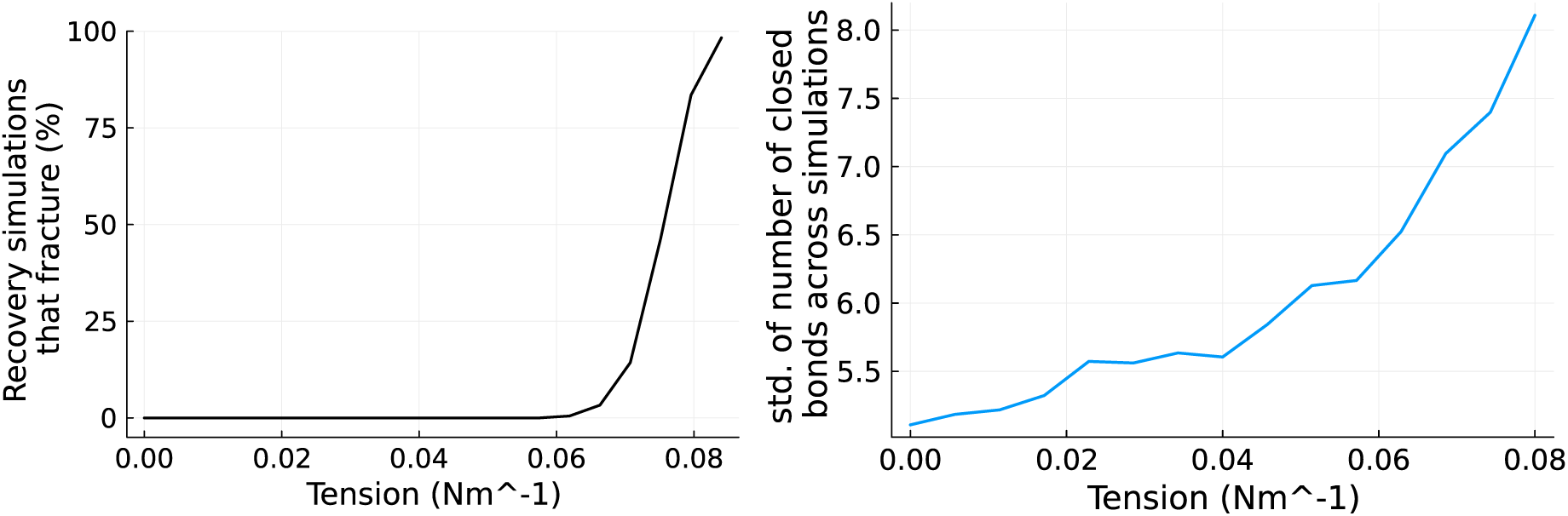
Effect of applied tension on repair timescale response. Left: percentage of recovery simulations resulting in rupture increases sharply close to Γ_c_, indicating increased noise close to the transition. Right: standard deviation of the number of closed bonds across simulations increases close to Γ_c_, also reflecting the increased variability in bond dynamics at the transition. Total simulation duration is 10,000s. Note: if the simulation duration were extended to infinity, all trajectories would eventually rupture due to stochastic fluctuation.

## 8 Alternative repair timescale, t_e_, stochastic model definition

**Figure 15:**
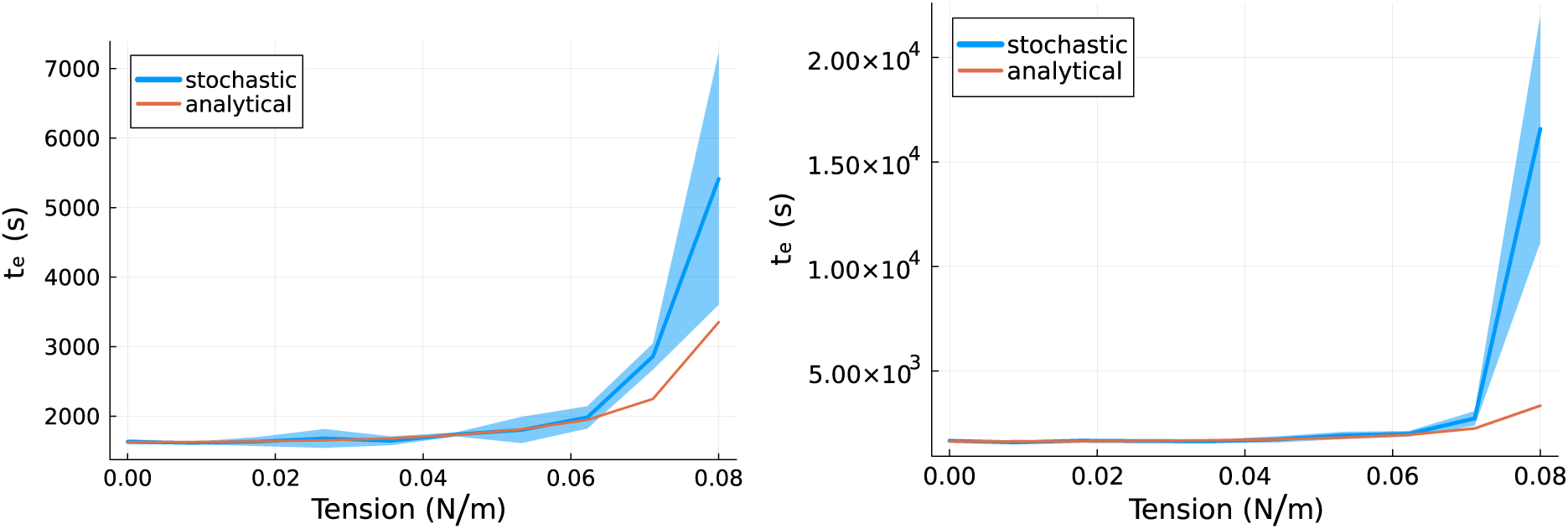
Alternative definition of the repair timescale, t_e_, for the stochastic model, where any trajectories that ruptured as the applied tension approached Γ_c_ were assigned an infinite rupture time and included only up to failure to capture initial recovery behaviour. The diverginf behaviourof the repair timescale close to Γ_c_ is now observed for both the analytical and stochastic models. Left: total duration 10,000. Right: total duration 20,000. The sharp divergence in the right plot suggests that, if simulated for sufficienctly long times, a system with a finite number of bonds will always eventually rupture.

## 9 Alternative initialisation strategy

**Figure 16:**
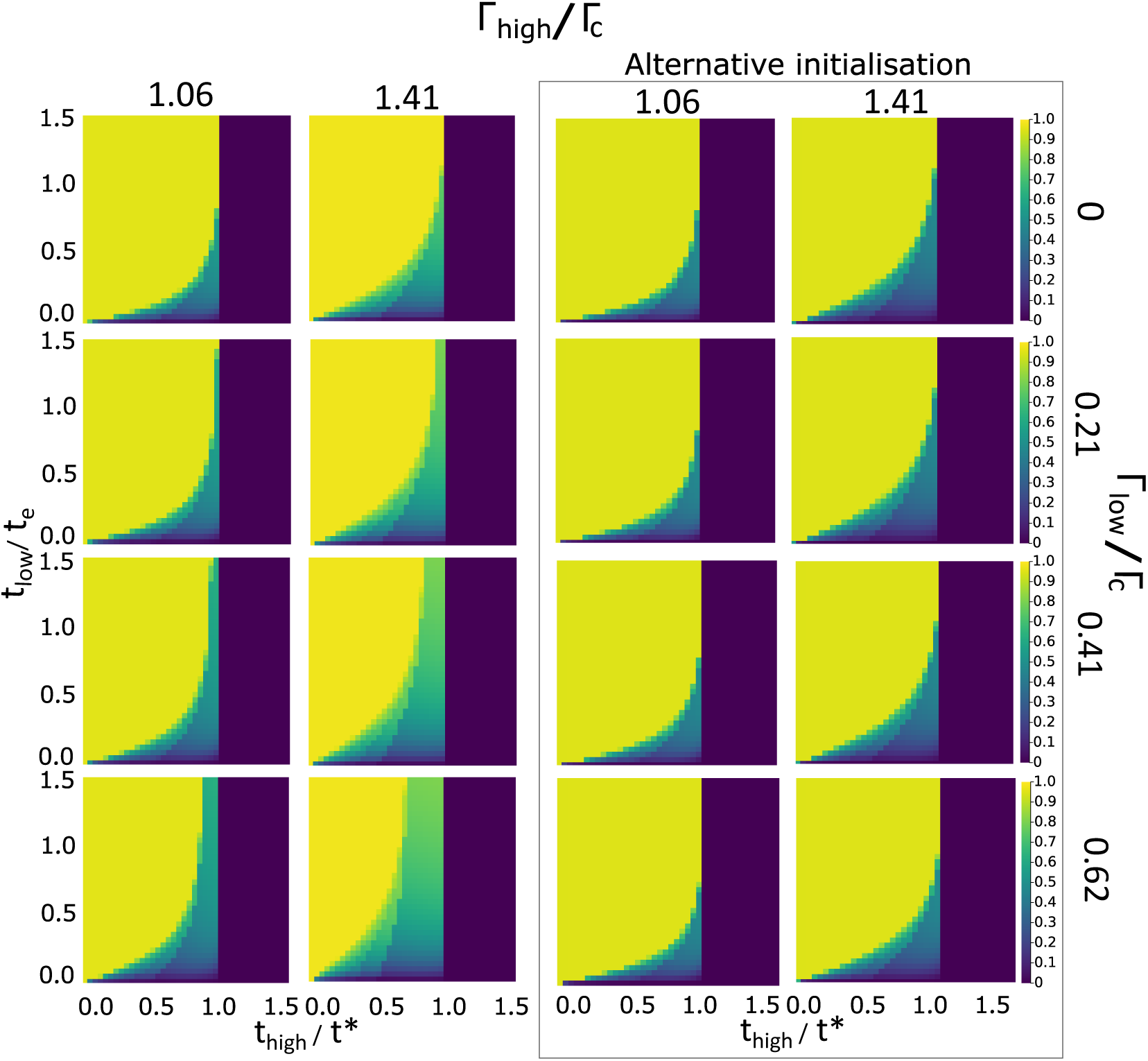
An alternative intitialisation strategy was considered for the map generation, whereby the initial number of closed links was set to the stable ratio of the specific low tension value (n_stable_(Γ_low_)). This removed the fourth regime of continuous diverging at both Γ_high_ and Γ_low_, as shown in the analytical model plots within the grey border.

## 10 Effect of doubling kinetic parameters while maintaining the k_on_/k_off_ ratio

**Figure 17:**
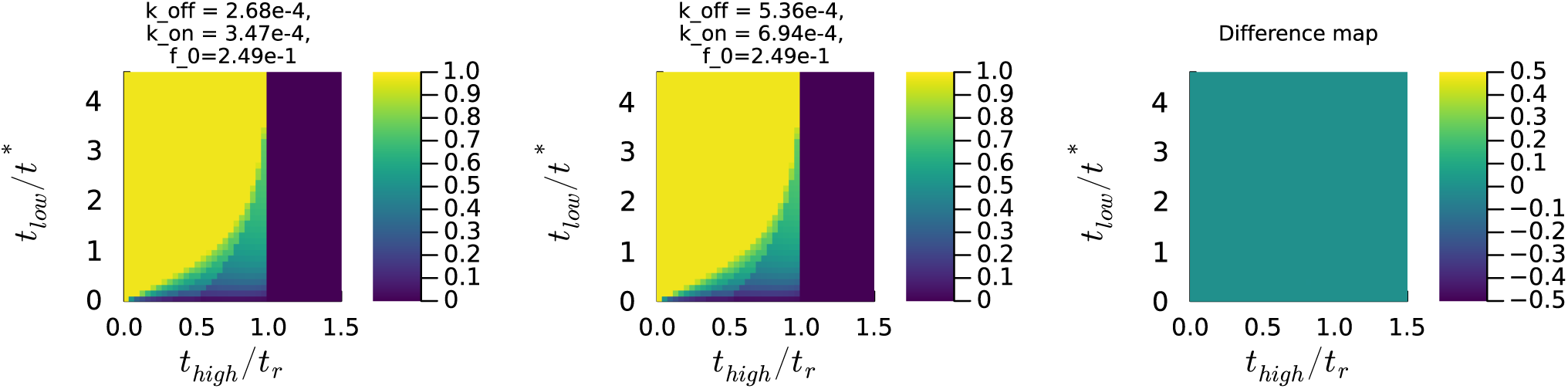
Heat maps show the normalised rupture probability as a function of the loading and relaxation durations (t_low_/t_e_ and t_high_/t^∗^) for the original parameter set (left) and for parameters doubled in magnitude while preserving the k_on_/k_off_ ratio (middle). The difference map (right) indicates negligible change across the parameter space, demonstrating that the model behaviour depends primarily on the relative balance between association and dissociation rates, rather than their absolute values.

## 11 Molecular basis of junction resilience: phase diagrams

**Figure 18:**
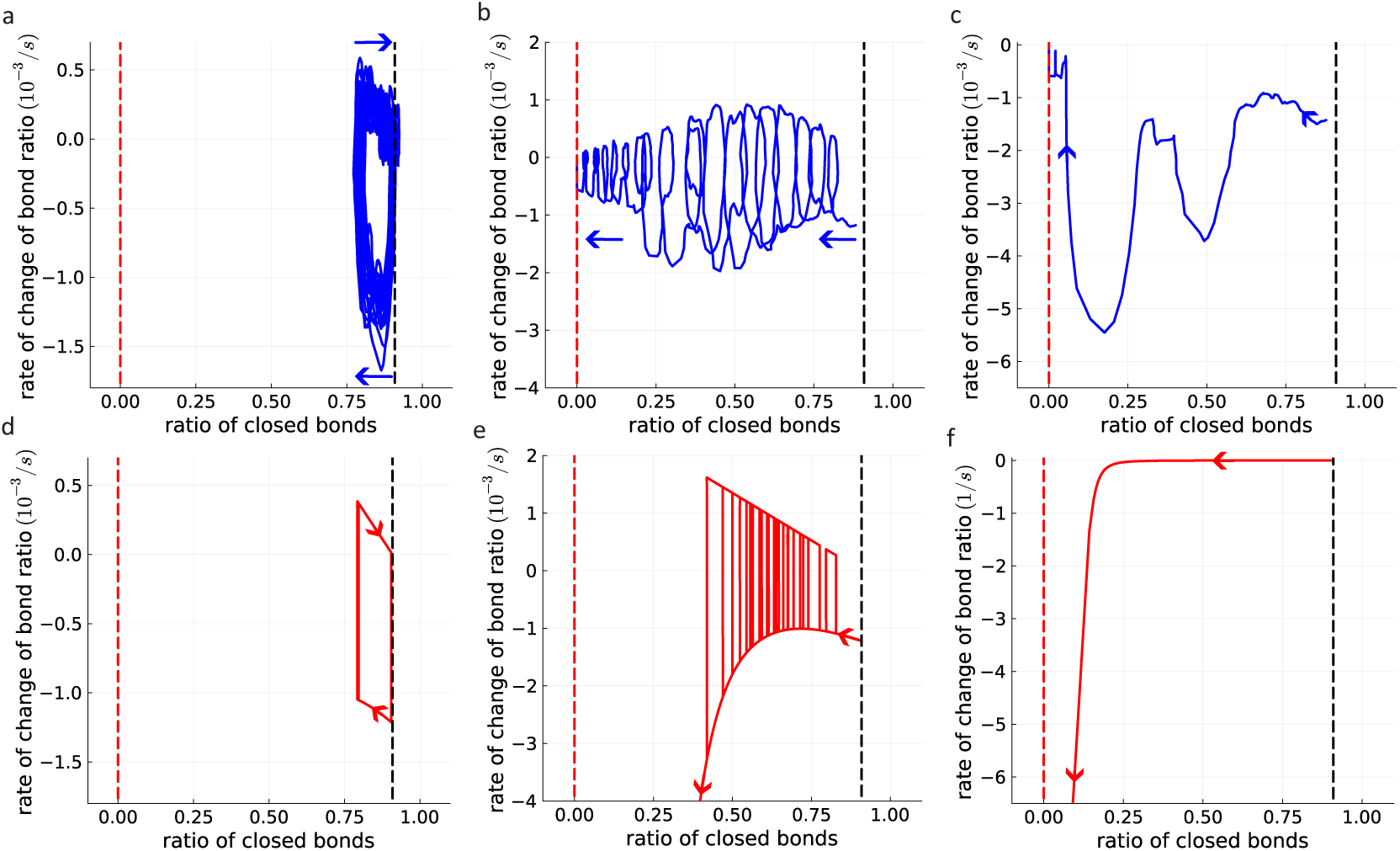
Phase-space trajectories of junctional bond dynamics under cyclic loading. Phase diagrams showing the rate of change of the fraction of closed intercellular bonds as a function of the instantaneous fraction of closed bonds for different combinations of high- and low-tension phases. Panels (a–c) show stochastic simulations of a single junction (blue), and panels (d–f) show corresponding analytical results (red). The black dashed lines mark stable fixed points, while red dashed lines denote unstable fixed points separating rupture and recovery regimes. Arrows indicate the direction of trajectory evolution during cyclic loading. Oscillatory trajectories (a,b) correspond to partial bond recovery within the slow-damage regime, whereas monotonic collapse to the left (c,f) reflects irreversible rupture. Stable limit cycles near the right boundary (a,d) indicate sustained recovery and junctional stability.

## 12 Maps scaled by rupture cycles (number of cycles-to-rupture)

**Figure 19:**
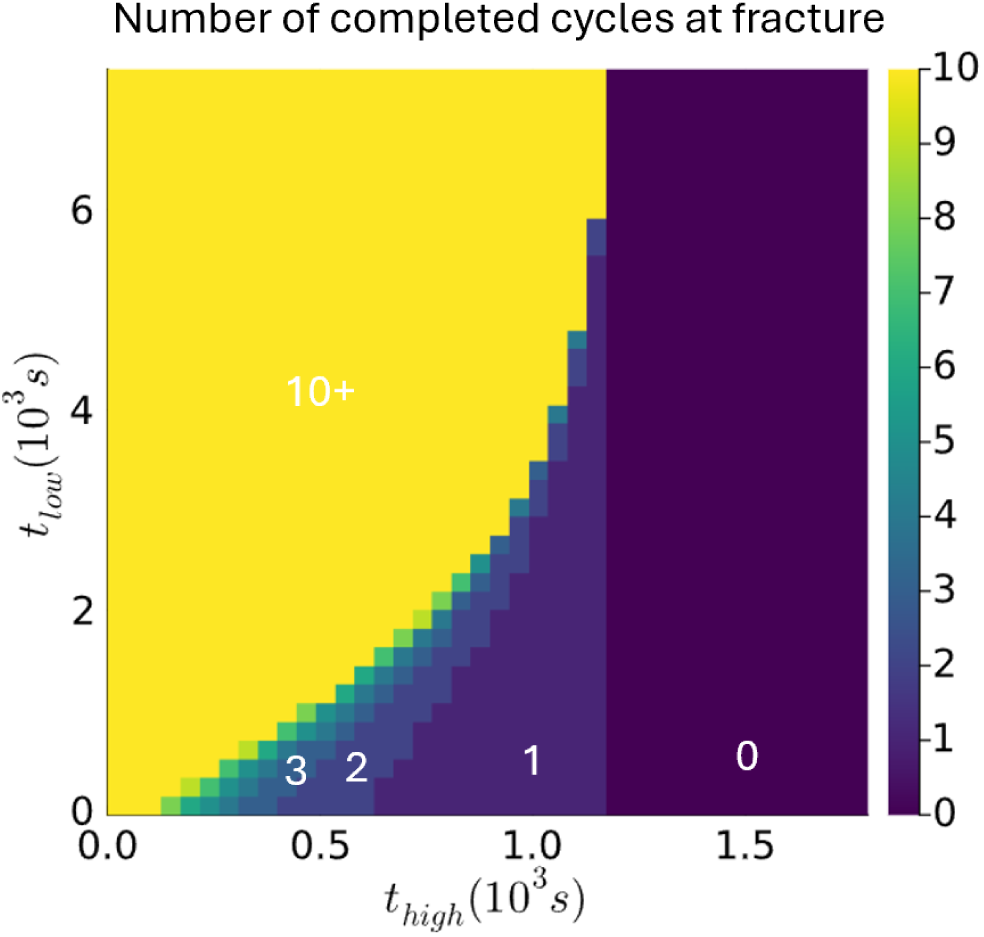
Number of completed load cycles at rupture as a function of high- and low-load durations. Colours indicate the number of cycles, with yellow representing 10 or more cycles (maximum duration) and purple indicating zero cycles (immediate rupture). In the slow damage accumulation regime, the fringes illustrate transitions to larger cycle numbers, in a direction towards smaller t_high_ and larger t_low_ indicating that slower damage accumulation and increased recovery extend the junction’s lifetime to a higher number of cycles.

